# Variable Number Tandem Repeats mediate the expression of proximal genes

**DOI:** 10.1101/2020.05.25.114082

**Authors:** Mehrdad Bakhtiari, Jonghun Park, Yuan-Chun Ding, Sharona Shleizer-Burko, Susan L. Neuhausen, Bjarni V. Halldórsson, Kári Stefánsson, Melissa Gymrek, Vineet Bafna

## Abstract

Variable Number Tandem Repeats (VNTRs) account for a significant amount of human genetic variation. VNTRs have been implicated in both Mendelian and Complex disorders, but are largely ignored by whole genome analysis pipelines due to the complexity of genotyping and the computational expense. We describe adVNTR-NN, a method that uses shallow neural networks for fast read recruitment. On 55X whole genome data, adVNTR-NN genotyped each VNTR in less than 18 cpu-seconds, while maintaining 100% accuracy on 76% of VNTRs.

We used adVNTR-NN to genotype 10,264 VNTRs in 652 individuals from the GTEx project and associated VNTR length with gene expression in 46 tissues. We identified 163 ‘eVNTR’ loci that were significantly associated with gene expression. Of the 22 eVNTRs in blood where independent data was available, 21 (95%) were replicated in terms of significance and direction of association. 49% of the eVNTR loci showed a strong and likely causal impact on the expression of genes and 80% had maximum effect size at least 0.3. The impacted genes have important role in complex phenotypes including Alzheimer’s, obesity and familial cancers. Our results point to the importance of studying VNTRs for understanding the genetic basis of complex diseases.

## 1 Introduction

The human genome consists of millions of tandem repeats (TRs) of short nucleotide sequences. These are often termed as Short Tandem Repeats (STRs) if the repeating unit is < 6bp, and Variable Number Tandem Repeats (VNTRs) otherwise. Together, they represent one of the largest sources of polymorphisms in humans^1–3^. While multiple resources have been developed for genome-wide analysis of STRs, here we focus specifically on VNTRs, which have been largely missing from genome-wide studies due to technical challenges of genotyping and the computational expense.

We define VNTR genotyping in the narrower sense of determining VNTR length (number of repeating units). As VNTRs can be located in coding regions^4^, untranslated regions^5^, and regulatory regions proximal to a gene^6, 7^, the variation in length can have an outsized functional impact. Not surprisingly, VNTRs have been implicated in a large number of Mendelian diseases that affect millions of people world-wide^8–10^. They also are known to modulate quantitative phenotypes in several other organisms^11^, and have shown pathogenic effects in other vertebrates^12, 13^. VNTRs have influenced the evolution of primates^14^ and specifically contributed to human evolution through gene regulation and differentiation of the great ape population^15^. Recent studies have identified VNTRs that have expanded in the human lineage or are differentially spliced or expressed between human and chimpanzee brains^16^.

Single nucleotide polymorphisms (SNPs) that associate with gene expression, often referred to as expression Quantitative Trait Loci (eQTLs), are molecular intermediates that drive disease and variation in complex traits^17–19^. Studies have shown that causal variants for diseases often overlap with cis-eQTL variants in the affected tissue^20, 21^. Therefore, we focus on the specific application of identifying expression mediating VNTRs (‘eVNTRs’), or VNTRs located in regulatory regions whose length is correlated with the expression of a proximal gene. Examples of ‘eVNTRs’ are: a) a VNTR in the 5’ UTR of AS3MT which is strongly associated with AS3MT gene expression and lies in a schizophrenia associated locus^5^; and b) a 12-mer expansion upstream of the cystatin B (CSTB) gene is associated with gene expression and with progressive myoclonus epilepsy^10, 22^.

Despite their importance, the full extent of VNTRs in mediating Mendelian and complex phenotypes is not known due to genotyping challenges. Traditionally, VNTR genotyping used capillary electrophoresis which did not scale to large cohorts. Despite the advent of sequence based genotyping, repetitive sequences continue to be challenging for genomic analysis^23^. For example, ‘stutter errors’ due to polymerase slippage during PCR amplification change VNTR length and reduce genotyping accuracy^24^. While tools for genotyping STRs have been developed^1,25,26^, they generally do not detect or genotype VNTRs, which have non-identical and larger repeat units. Recently, a few specialized computational methods (including our own) have been published to tackle the problem of genotyping VNTRs from sequence data^27, 28^. However, these methods are too computationally intensive to scale to functional studies with hundreds of individuals and 10^4^ VNTR loci (Results).

For these reasons, large-scale studies of VNTRs and their association with gene expression have been limited when compared to other sources of human variation such as SNPs and CNVs^21,29–31^. While the standard whole genome sequencing (WGS) frameworks often ignore repetitive regions^23^,there is some progress towards ‘harder’ variant classes such as eSTRs^32–34^ and ‘eSVs’^31, 35^. Therefore, ‘missing heritability’–the gap between estimates of heritability, measured for example by twin studies^36, 37^, and phenotypic variation explained by genomic variation– remains a limitation for eQTL studies^38^. It has been speculated that the inclusion of tandem repeats in association analyses may reduce this heritability gap^8,38,39^.

Here, we describe adVNTR-NN, a method that uses shallow neural networks for fast read recruitment followed by sensitive Hidden Markov Models for genotyping. We tested the speed and accuracy of adVNTR-NN on extensive simulations to demonstrate accuracy. We used adVNTR-NN to genotype over 10,000 VNTRs in 652 individuals from the GTEx project and associate VNTR length with gene expression in 46 tissues. We additionally validated eVNTRs in blood tissues in 903 samples from an Icelandic cohort and 462 samples from the 1000 genome project with Gene expression data (Geuvadis cohort). We compared the strength of eVNTR association against proximal SNPs to understand causality, and tested association with complex phenotypes. Our results suggest that it is computationally feasible to genotype VNTRs accurately in thousands of individuals, and multiple eVNTRs are likely to causally impact the expression of key genes involved in common and complex diseases.

## 2 Results

### Target VNTR Loci

Using Tandem Repeat Finder^40^, 502,491 VNTRs were identified in the GRCh38 human assembly. Over 80% of these had total length < 140bp (Fig. 1a) and could be genotyped using Illumina sequencing. As genotyping VNTRs remains computationally expensive, we focused on the 13, 081 VNTRs located within coding, untranslated, or promoter regions of genes (Methods) as they are most likely to be involved in gene regulation. Of those, we identified 10, 262 VNTRs that were within the size range for short-read genotyping (Fig. 1a). We added two additional VNTRs that were previously linked to a human disease (Supp Table S1) to obtain 10, 264 target loci^41, 42^.

**Figure 1:**
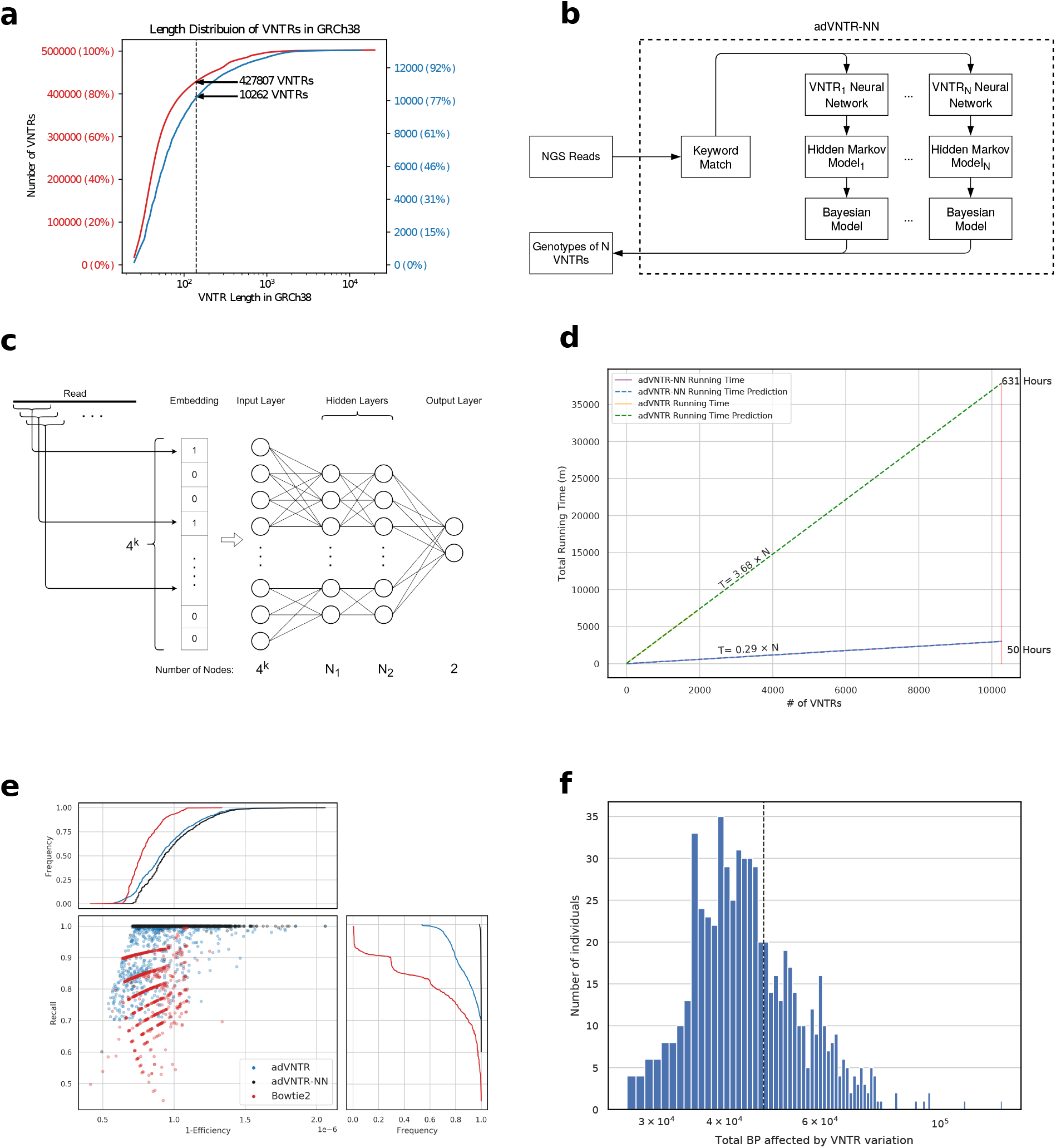
VNTR genotyping accuracy and speed. (a) Length distribution of all known VNTRs (red) and selected targeted VNTRs (blue) across the GRCh38 human genome. (b) The genotyping pipeline. (c) Neural network architecture for each VNTR which uses a mapping of reads to a k-mer composition vector. (d) Improvement in running time after using neural network and kmer matching. (e) Accuracy and efficiency of read recruitment. The scatter plot shows 1-efficiency ((TP+FP)/R) and recall (TP/(TP+FN)) of classification with different methods. High efficiency is related directly with running time. Each of 10,264 points represents a VNTR locus (method) and are shown once for each method. The side and top panels show cumulative distributions of recall and 1-efficiency. (f) Base-pairs (log-scale) affected by VNTRs per individual.

### 2.1 adVNTR-NN improves genotyping speed

Our previously published tool, adVNTR, used customized Hidden Markov Models (HMMs) for each VNTR and showed excellent genotyping accuracy, based on trio-analysis, simulations and PCR^27^. However, HMMs are compute-intensive, and despite some filtering strategies used by adVNTR(Methods), the time to genotype n=10K VNTRs was about 631 hours per individual. In developing adVNTR-NN, we first made significant improvements to pre-processing time. Next, we deployed a second filtering step with a 2-layer feed-forward network trained separately for each VNTR that accepted the k-mer composition for each read and filtered it specifically for that VNTR (Fig. 1b,c and Methods). The neural-network filter required 0.03s per read, and filtered reads with high efficiency in filtering reads. For 55X whole genome sequencing (WGS) with *r* = 4.2 × 10^6^ unmapped reads, the NN supplied an average of 14 previously unmapped reads to each VNTR HMM. Combining with the mapped reads, each HMM received an average of 32 reads per VNTR locus. This reduced the running time for *n* VNTR loci

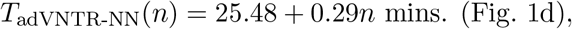

to allowing each individual to be genotyped at n=10K VNTRs in 50 CPU hours, a 13× speedup over adVNTR.

### adVNTR-NN outperforms alternative alignment strategies at VNTRs

While adVNTR was highly accurate by itself, its final accuracy depended upon reads filtered for genotyping, and specifically on false negatives–reads that were incorrectly removed by a filter. Formally, a read sampled from a VNTR was considered to be true positive (TP) if it passed the filter for that VNTR, and false negative (FN) otherwise. False positives (FP)–reads that passed the filter despite not being from the VNTR locus–were a lesser concern because they would eventually be discarded by the HMM for not aligning well to the model. However, high false-positives increase the running time. To account for this, we measured the trade-off between efficiency (1 −(TP+FP)/r) and recall TP/(TP+FN).

For comparisons with alternative filters, we used Bowtie2 as a representative read-mapping tool^43^. These tools are designed for fast mapping of reads and are accurate for most of the genome, but are not specifically designed for VNTR mapping genotyping (could have high FN). As a second comparison, we used adVNTR^27^, which has high recall (low FN), for VNTR mapping, and other graph based models in terms of sensitivity^44, 45^. We used a mix of real and simulated reads to test performance (Methods).

In terms of efficiency (1-(TP+FP)/r), Bowtie2 was the most efficient retaining only 0.9 in 10^6^ reads for further processing for 90% of the VNTRs. Both adVNTR and adVNTR-NN were slightly less efficient retaining about 1.2 reads per million for 90% of the VNTRs. However, they had significantly better recall. adVNTR-NN filtered reads with at least 90% recall for 99% of the target VNTR loci (Fig. 1e). In comparison, 80% of the loci achieved that recall for adVNTR, and only 27% of the loci had a recall of 90% for Bowtie2. Notably, adVNTR-NN had much better recall compared to adVNTR while also being more efficient, and therefore faster.

We had previously shown that improvement in recruitment improves genotyping accuracy^27^. Here, we used a mix of whole genome sequencing data and simulated reads (Methods) to compare the overall running time and accuracy of adVNTR-NN genotyping with VNTRseek^28^, which was not available at the time of original release of adVNTR. Notably, VNTRseek combines VNTR discovery and genotyping and does not customize genotyping for each VNTR. Therefore, its running time on 55X WGS ranged from 9640-9686 minutes, and was largely independent of the number of target VNTRs (Supp. Fig. S1). This was in contrast to the 1,696 minutes required by adVNTR-NN. The speed advantage for adVNTR-NN could largely be attributed to filtering strategies which could potentially be used to improve VNTRseek genotyping time as well. On simulated heterozygous reads with 30X coverage (Methods), adVNTR-NN was highly accurate. It achieved 100% accuracy in 7343 (76%) of 9638 VNTRs compared to VNTRseek’s median accuracy of 60% (Supp. Fig. S3). In contrast with adVNTR-NN, VNTRseek’s genotyping accuracy was sharply asymmetric, with much lower accuracy for decreasing VNTR length (Supp. Fig. S2).

### 2.2 Profiling eVNTRs

#### Data

To identify expression-mediating VNTR Loci (*eVNTRs*), we primarily used data from the GTEx project^21^ (Methods). The GTEx project provided WGS for 652 individuals as well as RNA-seq for each of these individuals from 46 tissue types including whole-blood. A majority (86.0%) of the donors were of European origin; another 11.5% were African American and the remaining were Asian and American Indian. For validation, we used a second cohort of 903 *Icelandic* individuals^46^ with associated whole blood RNA expression data and WGS. We also chose a smaller, third cohort from the *Geuvadis*^30^ project which provided gene-expression data in lymphoblastoid cell-lines for 462 samples, where the WGS for the samples was available from the 1000 genomes project^47^. 80.7% of the Geuvadis cohort was individuals with predominantly European (80.7%) ancestry and the remaining had African ancestry (19.3%). Due to the match of tissue type and ethnicity, the Icelandic and Geuvadis whole blood data were used for validation of methods for identifying eVNTRs discovered from the GTEx project.

#### eVNTR identification

We genotyped 10,264 VNTR loci in all 652 samples from GTEx to study the role of VNTRs in mediating gene expression of proximal genes. As expected, the most frequent allele matched the reference allele in 96.8% of the cases (Supp. Fig. S4).

Despite the GTEx data being predominatly European, 51% of the target VNTRs were polymorphic. Consistent with evolutionary constraints, VNTRs in promoters were most likely to be polymorphic (57%) followed by Untranslated regions (UTRs) (51%) and coding exons (47%) (Fig. 2a). Each individual in the GTEx cohort had a non-reference allele in at least 839 of the tested VNTR loci, with an average of 1,259 non-reference VNTRs per individual. Altogether, the 10,264 VNTRs inserted or deleted an average of 47,197bp per individual (Fig. 1f). As this represents < 10% of all VNTRs, the results highlight VNTRs as an important source of genomic variation. The minimum variation in a non-reference VNTR allele involved at least 6 basepairs and the average change in each variant site was 37bp or about 3 repeat units (Suppl. Fig. S5).

**Figure 2:**
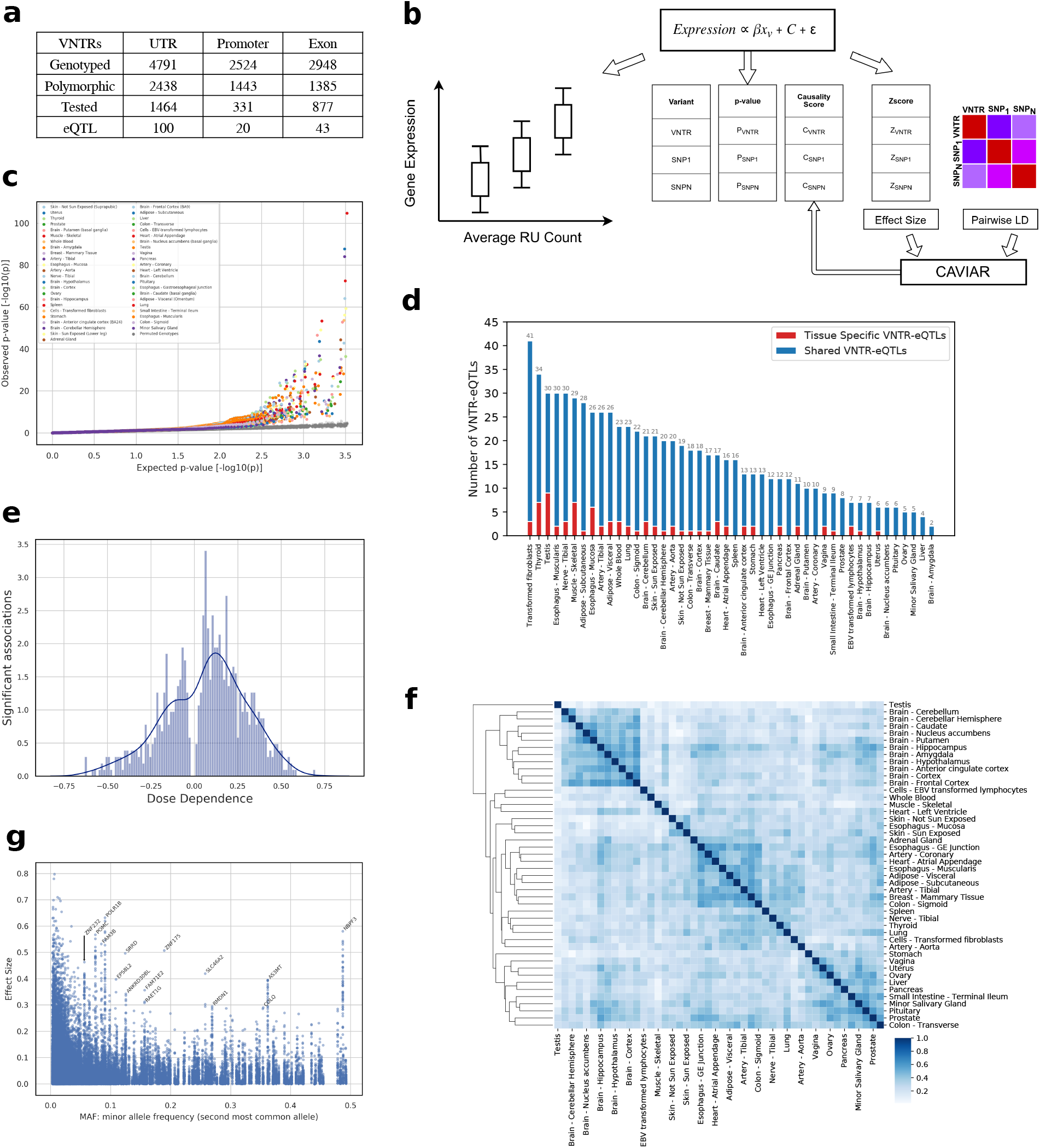
Effect of VNTR genotypes on mediating gene expression. (a) Location of target VNTRs and e-VNTRs relative to the proximal genes. (b) Pipeline to identify eVNTRs and assign causality scores. (c) Quantile-quantile plot showing p-values of association signals separated by tissue. Green line represents the p-values of 1,000 permutations. (d) Number of unique and shared eVNTRs in each tissue. (e) Trend of RU count correlation with gene expression level. (f) Spearman correlation of eVNTRs effect sizes for each pair of tissues. (g) Scatter-plot correlating effect size versus Minor Allele Frequency (MAF).

We excluded VNTRs that were monomorphic (1817), violated Hardy-Weinberg equilibrium constraints (1445) or had low minor allele frequency (<1%) after removing individuals in the GTEx cohort with no expression data for the specific gene (4330) (Methods), resulting in 2,672 genotyped VNTRs for association analysis. We used linear regression to measure the strength of association between average VNTR length of the two haplotypes, and adjusted gene expression level of the closest gene (Fig. 2b and Methods). To account for confounding factors, we included sex and population principal components of each individual as covariates. We also added PEER (probabilistic estimation of expression residuals) factors to account for experimental variations in measuring RNA expression levels (e.g batch effects, environmental variables)^48^. Briefly, PEER infers hidden covariates influencing gene expression levels, and we removed their effect by producing a residual gene expression matrix and using it for linear regression (See Methods).

We measured association with gene expression in each of the 46 tissues. To control False Discovery Rate (FDR), we used the Benjamini-Hochberg procedure to identify a tissue-specific 5% FDR cutoff (Supp. Fig. S6 and Methods). Combining data from all tissues, 759 tests tied to 163 unique VNTR loci passed the significance threshold (Fig. 2c). We refer to these (VNTR, gene) pairs as eVNTRs. The strength of association did not depend upon the location of the VNTRs in either core promoter, UTR, or coding regions. (Supp. Fig. S7). However, we VNTRs within 100bp of the Transcription Start Sites (TSS) were twice as likely to be eVNTRs compared to other locations (P = 6 × 10^−6^; Fisher’s exact test), consistent with their known roles in core-promoters^49^.

The number of eVNTRs observed in each tissue type was different but mostly consistent with the number of individuals samples for each tissue type. Only 4% of the eVNTRs were tissue specific, with each tissue containing a similar number of tissue specific eVNTRs (Fig 2d). An analysis using mash^50^ showed that many (38%) eVNTRs were significant in at least half (23) of the tissues tested (Fig. S8).

Twenty-three of the 163 unique eVNTRs showed significant association in whole blood (Table 1), a tissue type in which we could validate the eVNTRs using independent data from the Icelandic cohort of 903 individuals. Two of the 23 VNTR loci could not be genotyped in the Icelandic cohort due to missing data. 18 (86%) of the 21 VNTRs showed significance at a similar level and same direction of effect in Icelanders, highlighting the strong reproducibility of the associations. The Geuvadis data were acquired for a smaller cohort compared to the Icelandic data and measured expression in lymphoblastoid cells–transformed B cells, which are a component of whole blood tissue. Nevertheless, 12 of the eVNTRs were replicated. Combined, 91% (20/22) of eVNTRs could be replicated in an independent cohort where data was available.

**Table 1:** Replication of whole blood VNTRs in independent cohorts. Each row describes an eVNTR in whole blood from GTEx project(n=652 individuals) identified with false discovery rate (FDR) < 0.05. Replication of the signal in whole blood tissue of the Icelandic cohort of 903 samples and in lymphoblastoid cell-lines from the Geuvadis cohort (462 samples) with the same direction of effect and FDR < 0.05. Length (respectively, RU length) refers to the total (respectively, repeat-unit length) of the VNTR.

|  |  |  |  |  |  |  | Validation |  |
| --- | --- | --- | --- | --- | --- | --- | --- | --- |
|  | Locus | Length | RU Length | Effect Size | Gene | Annotation | Icelandic | Geuvadis |
| 1 | chr1:21440112-21440147 | 35 | 6 | 0.43 | NBPF3 | UTR | Y | Y |
| 2 | chr2:24084339-24084414 | 75 | 25 | -0.12 | TP53I3 | UTR | Y | Y |
| 3 | chr2:25161573-25161616 | 43 | 9 | 0.22 | POMC | Coding | Y | Y |
| 4 | chr2:112542424-112542500 | 76 | 25 | -0.18 | POLR1B | Coding | Y | Y |
| 5 | chr3:56557249-56557289 | 40 | 20 | -0.12 | CCDC66 | Coding | Y | Y |
| 6 | chr6:13328502-13328532 | 30 | 6 | 0.12 | TBC1D7 | UTR | Y | Y |
| 7 | chr7:64337190-64337240 | 50 | 13 | 0.09 | ZNF736 | UTR | Y | Y |
| 8 | chr8:86508719-86508765 | 46 | 23 | 0.13 | RMDN1 | UTR | Y | Y |
| 9 | chr10:102869497-102869605 | 108 | 36 | 0.22 | AS3MT | Coding | Y | Y |
| 10 | chr21:46228815-46228863 | 48 | 9 | -0.03 | LSS | UTR | Y | Y |
| 11 | chr17:75589192-75589228 | 36 | 6 | -0.06 | MYO15B | Coding | Y | - |
| 12 | chr1:46609102-46609134 | 32 | 16 | 0.09 | MOB3C | UTR | Y | N |
| 13 | chr5:80654880-80654954 | 74 | 9 | 0.04 | MSH3 | Coding | Y | N |
| 14 | chr9:137063433-137063550 | 117 | 39 | -0.15 | SAPCD2 | UTR | Y | N |
| 15 | chr14:61762420-61762454 | 34 | 17 | 0.03 | SNAPC1 | UTR | Y | N |
| 16 | chr19:12577507-12577551 | 44 | 22 | -0.09 | ZNF490 | UTR | Y | N |
| 17 | chr21:41316673-41316756 | 83 | 13 | -0.19 | FAM3B | UTR | Y | N |
| 18 | chr22:37805258-37805313 | 55 | 6 | 0.11 | H1F0 | UTR | Y | N |
| 19 | chr1:202187007-202187042 | 35 | 7 | 0.06 | PTPRVP | UTR | N | Y |
| 20 | chr17:18208488-18208544 | 56 | 7 | -0.13 | ALKBH5 | UTR | N | Y |
| 21 | chr17:76564106-76564152 | 46 | 9 | 0.11 | SNHG16 | UTR | - | N |
| 22 | chr17:56978047-56978107 | 60 | 20 | 0.15 | SCPEP1 | UTR | N | N |
| 23 | chr6:30163542-30163579 | 37 | 12 | 0.14 | TRIM15 | UTR | - | - |

In 65% of the cases, VNTR length had a positive correlation with gene expression; the remaining cases had a negative correlation (Fig. 2e). This was consistent with the hypothesis that many VNTRs encode transcription factor binding sites and increasing length improved the TF binding affinity. Moreover, the overall effect size was also large and 80% of the eVNTRs had a maximum effect-size 0.3 or higher.

We computed correlation of eVNTR effect size between each pair of tissues using the Spearman rank test. Despite the multi-tissue activity of most eVNTRs, each tissue showed distinct behavior with low correlation to most other tissues (Fig. 2f). Similar tissue types were expectedly correlated (e.g. brain). Some correlations were seen among glandular tissues (salivary, prostate, pitutary) and also between adipose tissue and nearby tissues and organs (heart, esophagus, artery, breast). Thus, even though most eVNTRs are shared across tissues, we hypothesize that the combined effect of active eVNTRs is tissue-specific and leads to unique regulatory program for each tissue type.

Similar to SNPs, VNTR loci generally showed a negative correlation between Minor Allele Frequency (MAF) and effect size, so that common variants generally had low effect size with larger effects mainly shown by rare variants^51^ (Fig 2g). However, we still observed many eVNTRs where common VNTR (MAF > 0.05) showed large effects. These eVNTRs had highly significant p-values (Supp. Fig. S9) and in many cases, the proximal genes were associated with known diseases or phenotypes (Table 2). As these represent potentially the most interesting eVNTR findings, we tested them further for causality and function.

**Table 2:**
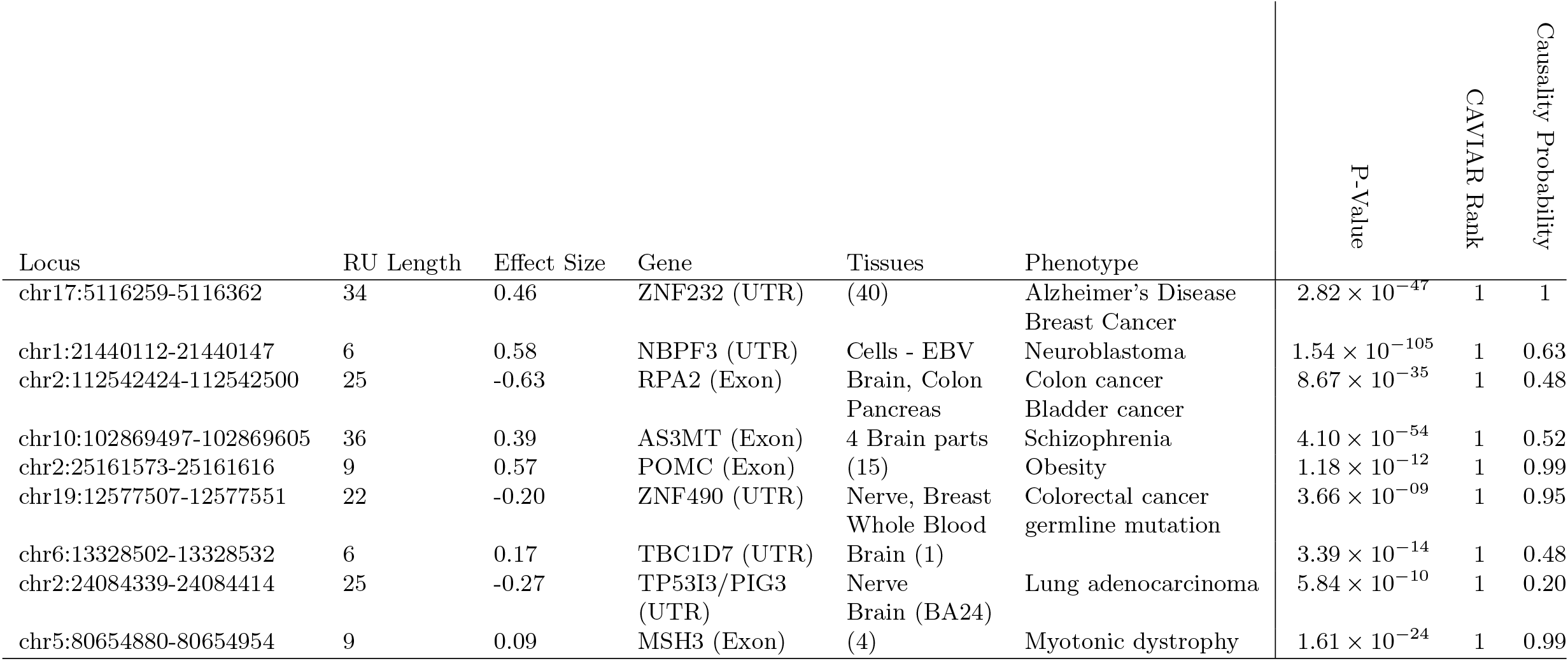
A partial list of genic eVNTRs in GTEx cohort with known phenotypes of the related genes. Top tissues are noted except in cases where significance is seen in 4 or more tissues.

#### VNTRs mediate expression of key genes

Only a small number of examples have been reported where VNTR repeat unit counts have a causative on gene expression^5^. One well known example is the AS3MT gene which is involved in early brain development, where the VNTR was associated with expression and was in LD with SNPs associating with schizophrenia^5^.

To investigate causality, we ranked each eVNTR against all SNPs within 100kbp by (a) comparing the relative significance of association with gene expression; and (b) using the tool CAVIAR^52^ to measure the causality of association (Methods). Remarkably, the two rankings were very similar with mean discrepancy 2|*r*_1_ − *r*_2_|/(*r*_1_ + *r*_2_) = 2.3 × 10^−3^ across the 163 eVNTRs. We used the harmonic mean (2/ (1/*r*_1_ + 1/*r*_2_)) of the two ranks to order the eVNTRs. Of the 163 VNTRs, 111 had a harmonic rank ≤ 10 and 81 of the eVNTRs were ranked 1 (Supp. Fig. S10), indicating that the majority of the eVNTRs could be considered causal in some tissue. Separating tissue types, 170 (22%) of the 759 significant associations were likely causal. These results suggest a much larger causality fraction compared to SNPs, structural variants^31^ and even STRs^34^, even with the caveat that we only tested ‘genic’ VNTRs.

Looking at individual eVNTRs, we recapitulated a previous result by identifying an eVNTR in the AS3MT gene. The lowest association p-value measured in any tissue using 652 samples was 4.1 × 10^−54^, which was orders of magnitude higher than the significance reported with 322 samples^5^(Fig. 3a,b). Its harmonic rank for the two causality tests was 1. Finally, the VNTR is located in a regulatory region of the genome as identified by H3K27Ac and DNase marks (Fig. 3c).

**Figure 3:**
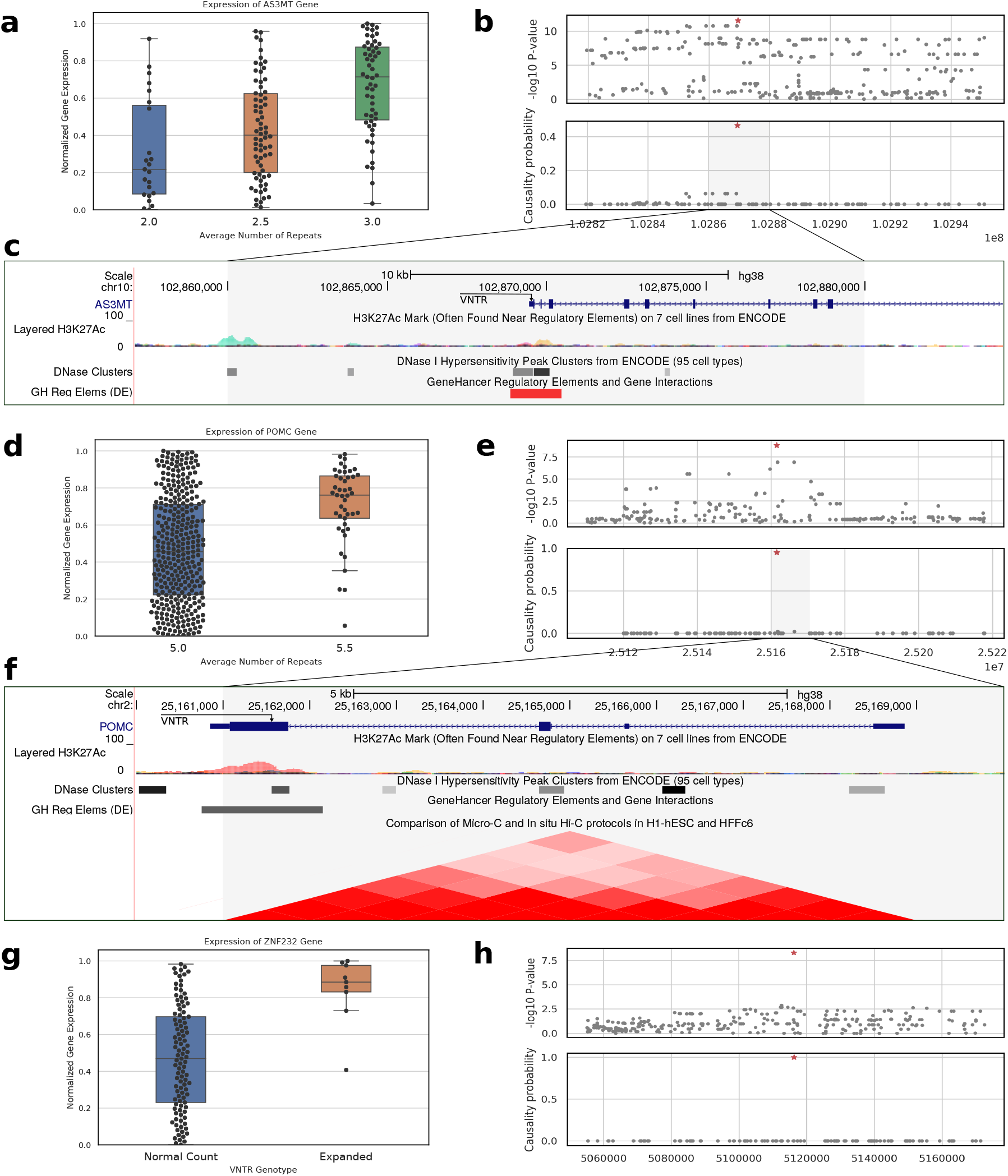
Effect of VNTR genotypes on mediating gene expression. (a) Association of AS3MT VNTR genotype with gene expression in Brain-Cortex (p-val:2.78 × 10^−12^). (b) Association with gene expression (upper panel) and CAVIAR causality probability of proximal SNPs– all SNPs in 100kbp window on either side of the AS3MT VNTR (red-star). (c) Location of AS3MT VNTR relative to known regulatory elements. (d,e): Association with gene expression of the POMC VNTR (p-val:1.53 × 10^−9^) and its causality probability relative to proximal SNPs. (f) Location of POMC VNTR relative to other regulatory regions and its spatial proximity with the promoter region revealed via Hi-C. (g,h) Association with gene expression of the ZNF232 VNTR (p-val:5.47 × 10^−9^) and its causality score relative to proximal SNPs.

Proopiomelanocortin (POMC) is a precursor of many peptide hormones with multiple roles including regulation of appetite and satiety^53^. Hypermethylation of POMC (and reduced expression) in peripheral blood cells and melanocyte-stimulating hormone positive neurons was strongly associated with obesity and body mass index^54, 55^. Surprisingly, POMC over-expression also predisposed lean rats into diet-induced obesity^56^. Our analysis identified a VNTR in the coding region of the POMC gene as the causal variant governing expression levels in 15 tissues, including adipose and nerve tissues. The 6R allele had 1.8-fold higher expression in blood and nerve cells (Fig 3d), and the correlation with expression was much stronger than neighboring SNPs (Fig 3e). Moreover, the VNTR was located within an H3K27Ac mark that was topologically close to the promoter of the gene based on chromatin conformation (Fig 3f).

The ZNF232 gene is differentially expressed in ovarian and breast cancers^57, 58^. Also, the chr17 locus containing the gene has been associated with Alzheimer’s in a recent large meta-GWAS study on the UK Biobank data^59^. We identified an eVNTR in the promoter region where expanded alleles (RU5+) had 2-fold higher median expression relative to RU3 (Fig. 3g). The VNTR was ranked 1 in 40 of 46 tissues including 7 brain sections, and specifically the Hippocampus, which is the affected region in Alzheimer’s^60, 61^ (Fig. 3h). It was also ranked 1 in ovary and breast with a normalized effect size that was twice the effect size of the best SNP (Table 2).

The RPA2 gene product is part of the Replication Protein A complex involved in DNA damage checkpointing^62^. Its over-expression is identified as a prognostic marker for colon cancer^63^ and bladder cancers^64^. A VNTR that overlapped the Transcription Start Site (TSS) of RP2A with lower VNTR length showed 1.9-fold higher expression of RPA2 in multiple tissues including colon (Supp. Fig. S11 and Table 2). Table 2 identifies other important genes including NBPF3 (Neuroblastoma^65^), TBC1D7 (lung cancer^66^), ZNF490 (colorectal cancer^67^), MSH3 (myotonic dystrophy^68^) and others. Taken together, our results suggest that VNTRs mediate the expression of key genes.

## 3 Discussion

VNTRs are the “hidden polymorphisms.” Despite high mutation rates and known examples of function modifications, VNTR analysis is not a component of Mendelian or GWAS analysis. This primarily is due to technical challenges in VNTR genotyping. Here, we use a combination of fast filtering followed by a hidden markov model-based genotyping to accurately determine VNTR genotypes. Our method can genotype 10K VNTRs for an individual in 50 hours making the time problem tractable. We used it to genotype close to 2, 000 human samples. The use of neural networks as a filtering strategy is novel, and we believe that further improvements could lead to another order of magnitude reduction in compute time, making it practical to genotype ≥ 10^5^ individuals in the future.

Some VNTRs have complex multi-repeat structure making it difficult to map reads and count the repeating units. However, unlike other VNTR genotyping methods, our method customizes the genotyping for each VNTR. Future research will focus on improving the genotyping for the hard cases, possibly by building HMMs with separate profiles for each distinct repeating unit, as well as the use of long-reads to improve anchoring to the correct locations. We pursue a targeted genotyping approach which has the disadvantage of not being able to discover new VNTRs, and we rely on other methods for the initial discovery of VNTRs. However, we note that the discovery is a one-time process while genotyping must be repeated for each cohort, and therefore, it makes sense to separate the two problems.

We found that VNTRs were strongly associated with the expression of proximal genes with over 6.1% of the tested VNTRs showing genome wide significant association. Importantly, nearly half of the eVNTR loci were more significant compared to neighboring SNPs, suggesting that a much higher fraction of eVNTRs are causal relative to other variant classes such as SNPs, structural variants, and even STRs. While the high fraction of causal eVNTRs can partly be explained by the choice of ‘genic’ VNTRs for testing, we note that it was computed only for eVNTRs, and speculate that eVNTRs identified in non-genic regions are located in regulatory regions and will continue to have stronger associations compared to neighboring SNPs. In summary, ongoing technical innovations in speed and accuracy of VNTR genotyping are likely to improve our understanding of human genetic variation, and provide novel insights into the function and regulation of key genes and complex phenotypes.

## Acknowledgements

The research was supported in part by grants HG010149, and R01GM114362 from the NIH. The analyses presented in this paper are based on the use of study data downloaded from the dbGaP web site, under phs001095.v1.p1, phs001096.v1.p1 and phs001097.v1.p1. This work used data from UK Biobank (project 46122). The 30X whole genome sequencing data of 1000 Genomes Project samples used in this research were generated at the New York Genome Center with funds provided by NHGRI Grant 3UM1HG008901-03S1.

## Conflict

V.B. is a co-founder, serves on the scientific advisory board, and has equity interest in Boundless Bio, inc. (BB) and Digital Proteomics, LLC (DP), and receives income from DP and BB. The terms of this arrangement have been reviewed and approved by the University of California, San Diego in accordance with its conflict of interest policies. BB and DP were not involved in the research presented here.

## 4 Method

### 4.1 Genotyping in adVNTR-NN

#### Filtering trade-off calculations

Let *A*(*r*) denote the HMM genotyping time using *r* reads. The goal of filtering is to reduce the number of reads supplied to each VNTR HMM. Any filter is characterized by three parameters:

**run-time:** Let *P* (*r*) denote the running time of the filter for *r* reads for each VNTR locus;
**efficiency:** Let *f_k_* denote the fraction of reads that were retained for any VNTR. The efficiency is defined as 1 − *f_k_* so that high efficiency implies only a small fraction being retained by the filter.
**sensitivity/recall:** The fraction of true VNTR overlapping reads that were accepted for each VNTR.

Consider a data-set with *r* unmapped reads and among the mapped reads, an average of *r*′ reads are assigned to each VNTR locus. Assuming that the filtered reads are distributed equally among the VNTRs, each HMM will receive *fk*_*r*_ + *r*′ reads on the average. The total genotyping time for *n* VNTRs is given by:

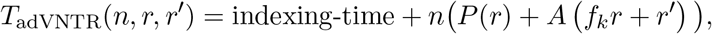

Empirically, *A*(*r*) = 0.32*r* seconds per VNTR. The keyword match filter for adVNTR achieved *f_k_* = 7.7 × 10^−5^. For a 55X coverage WGS with *r* = 4.2 × 10^6^ reads, *P* (*r*) = 111.22(*s*), *r*′ = 18, we run the HMM on an average of *fk*_*r*_ + *r*′ = 341 reads per VNTR on the average. The running time is:

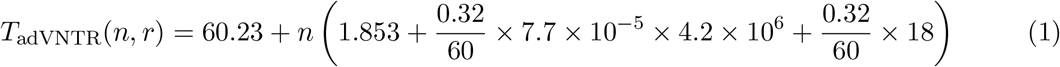

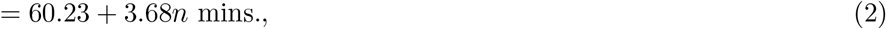

The genotyping time for n=10K VNTRs is about 631 hours per individual.

#### Read Filtering

For each VNTR locus *V*, and each read *R*, consider a binary classification function *f*: *V* × *R* → {0, 1}, where *f* (*R, V*) = 1 if and only if read *R* maps to locus *V*. For each read and each of *N* loci *V*_1_,…, *V_N_*, the neural recruitment method computes independent classification functions *f_i_*(*V_i_*, *R*). Note that a read can be assigned to multiple VNTR loci, or to none. As an initial step toward this task, we perform a fast string matching based on prefix tree (trie) to assign each read to the VNTR loci that share an exact match with the read. For an efficient matching, we generate a separate aho-corasick trie using every k-mer in VNTR loci as dictionary *X*. A trie is a rooted tree where each edge is labeled with a symbol and the string concatenation of the edge symbols on the path from the root to a leaf gives a unique word (k-mer) *X*. We label each leaf with a set of *T* VNTRs that contain corresponding k-mer. On the other hand, the string concatenation of the edge symbols from the root to a middle node gives a unique substring of X, called the string represented by the node. We add extra internal edges called failure edges to other branches of the trie that share a common prefix which allow fast transitions between failed string matches without the need for backtracking^80^. Testing whether a query *q* has an exact match in the trie can be done in *O*(|*q*|) and we require additional *O*(|*T* |) time to assign read *q* to all *T* VNTR loci that share the keyword. The overall complexity of this algorithm is linear based in the length of original dictionary (VNTRs in the database) to build the Trie and recover matches plus the length of queries (sequencing reads). Hence, after construction of the trie, the running time is proportional to just reading in the sequences.

#### Neural Recruitment

To further reduce the set of reads assigned to each VNTR, we use a 2-layer feedforward Neural Network to compute *f_i_*, using a *k*-mer based *embedding* to encode DNA strings. Specifically, we use a DNA string *w* of length *k*, consider an bijection *ϕ* that maps *w* to a unique number in [0, 4^*k*^ − 1]. Each read *R* can be defined by a collection of overlapping *k*-mers. We map read *R* to a unique vector *v_R_* ∈ {0, 1}^4*k*^, such that *v*_*R*_[*i*] = 1 if and only if *ϕ*^−1^(*i*) ∈ *R*. Deatils of the neural network architecture and hyper-parameters are presented below.

#### Network Architecture

Let *v* denote the mapping of a read. We use a shallow architecture with an input layer used to present *v* to the network. We add two layers of fully connected nodes as the hidden layers, with each node being a *Relu* function^72^. In the output layer, there are two nodes *zero* and *one* which specify that whether read should be classified as true (containing VNTR) or false (Fig. 1). We used the training set to train the network with Adam optimization algorithm^78^. The number of hidden layers *N*_1_ and *N*_2_ were chosen empirically. Too many nodes would increase both training time and test time and possibly cause over-fitting. We performed the training with the number hidden nodes of each layer varying from 10 to 100 with 10 increase in each step and selected *N*_1_ = 100 and *N*_2_ = 50 as the best parameters according to validation performance.

#### Choosing the optimal k-mer length

The choice of k-mer length is important. Increasing the k-mer size could decrease sensitivity in our case as small variation will significantly change the k-mer composition, whereas lowering k-mer size reduces the features that are discriminative for a pattern^70^. In addition, our embedding size exponentially grows with respect to the *k* so there is also a practical upper bound on the *k*. Following Zhang^70^ and Dubinkina^71^, we trained and tested in the range 4 ≤ *k* < 9. The accuracy remains comparable in this range (Fig. S12), and we chose *k* = 6 as its mean validation accuracy is the highest compared to four other values of *k*.

#### Effect of different loss functions

To choose the best loss function, we examined three regression loss functions: Mean Squared Error (MSE), Mean Squared Logarithmic Error (MSLE), and Mean Absolute Error (MAE), as well as three binary classification loss functions Hinge, Squared Hinge, and Binary Cross-Entropy. We compared the validation performance of our models for these 6 different loss functions. Each distribution in Supplementary Fig. S13 shows the accuracy on validation set across 1905 genomic loci. We analyzed these distributions using one-way analysis of variance (ANOVA) and none of them were significantly better than others. We chose binary cross-entropy as it obtained the highest mean accuracy (99.95%) among loss functions and its binary classification nature fits our requirement.

#### Speed and efficiency of neural network filtering

The neural-network filtering achieved a speed of *N* (*r*) ≃ 0.03*r* seconds for *r* reads, greatly increasing filtering efficiency (*f_n_f_k_*′ < 10^−6^) to input only 14 reads per VNTR on the average when *r* = 4.2 × 10^6^. The running time using the two filters could be modeled as

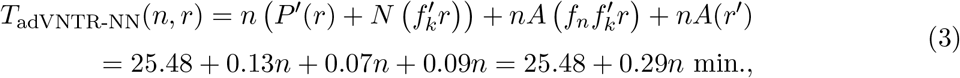

#### Simulated data for training and testing

We used ART^79^ to generate *r* = 6 × 10^8^ reads from human reference genome (30X coverage) with Illumina HiSeq 2500 error profile. For each target locus, we modified the number of the repeats to be ±3 of the original count in the reference with setting 1 as minimum number of repeats, and simulated reads from those regions. For each locus, we assigned labels to reads as being true reads or not, based on exact location. We divided the original set of reads into three parts: 70% for training, 10% for validation and 20% for testing. We trained all neural network models using the training and validation sets, and reported performance on the test dataset.

To augment the data, we added random single nucleotide variations in the genome sequences of the dataset^74^. For each sequence in the dataset, we replaced its nucleotides with a random one with probability *r_e_*. We set *r_e_* = 10^−5^, the novel base substitution mutation rate within VNTRs^81^. This method of dataset augmentation helps include ‘mutated’ k-mers in the embedding of reads, making the method more robust.

To test and compare genotyping accuracy against VNTRseek, we started with a random selection of 10,000 target VNTR loci (< 140 bp) and filtered them out if a VNTR locus was marked as indistinguishable in VNTRseek. As a result, 9,638 target VNTRs remained. We used ART^79^ to generate heterozygous samples by simulating 15X coverage reads from each modified haplotype which contained a non-reference allele and combined those with 15X reads that were simulated from reference. The non-reference allele for each VNTR was chosen to be in the range [*c* − 3, *c* + 3], where *c* is the reference count. Together, this provided six diploid simulated data-sets for each locus, at 30X coverage.

#### Performance test

We measured running time of adVNTR-NN and VNTRseek by running them with default parameters on a single core of Intel Xeon CPU E5-2643 v2 3.50GHz CPU. To measure the accuracy of genotyping, we ran adVNTR-NN and VNTRseek on diploid simulated data of heterozygous VNTRs and measured the number of correct calls divided by total number of VNTR loci.

### 4.2 Data and preprocessing

We accessed 30X Illumina WGS data from the GTEx cohort (652 individuals) through dbGaP (accession id phs000424.v8.p2). Specifically, we accessed CRAM files containing read alignments to the GRCh38 reference genome through cloud-hosted SRA data using fusera and downloaded VCF files containing SNP genotype calls from dbGaP.

As genotyping VNTRs remains computationally expensive, we focused on the smaller set of VNTRs located within coding, untranslated, or promoter regions of genes, which are most likely to be involved in regulation. We identified VNTRs in coding exons and UTRs by intersecting VNTR coordinates with refseq gene coordinates downloaded from UCSC Table Browser^86^. To identify VNTRs that appear within promoter regions, we considered 500bp upstream of the transcription start site of genes as the promoter regions. Overall, this procedure identified 13, 081 VNTRs, of which 10, 262 were within the size range for short-read genotyping (Fig. 1A). We subsequently added two VNTRs previously linked to a human disease to obtain 10, 264 target loci^42, 42^. We genotyped these VNTR loci in 652 individuals from GTEx cohort using adVNTR-NN on Amazon Web Services (AWS) cloud, which allowed us to do the computation in parallel for different samples.

We compared the most common allele of each VNTR with the reference allele (GRCh38) to observe representation of each VNTR in the reference. We also searched for VNTRs with multiple observed alleles to estimate a rate of polymorphism for VNTRs and find how common each allele was. To call a VNTR polymorphic, we set the minor allele frequency at 5% and any variation below that frequency was discarded. In addition, we identified the amount of base-pair difference that they make in genome of each individual by comparing the copy number difference of VNTRs between reference and the sample and multiplied that by the pattern length of each locus. We computed how many loci on average differed between an individual and reference by combining all non-reference calls in at least one haplotype from all individuals and dividing it by all called variants. VNTRs whose allele frequencies did not meet the expected percentage of homozygous versus heterozygous calls under HardyWeinberg equilibrium (*P* < 0.05 for two-sided binomial test) were eliminated. We further removed VNTRs that were monomorphic (only one allele) in the entire GTEx cohort or had minor allele frequency lower than 1% among the individuals with expression data in every tissue. We used the resulting 2,672 VNTRs for subsequent analysis (Supp Table S1).

We obtained processed RNA-expression data (RPKM values) from 54 tissues from dbGaP (phe000020.v1) and limited analysis to 46 tissues which had data for at least 100 individuals. ‘Non-expressed genes’– genes with median RPKM level zero– in each tissue were removed from analysis. For the remaining genes, we quantile-normalized RPKM values of each tissue to a normal distribution. We analyzed VNTR-Gene pairs for each VNTR and its closest gene based on refseq annotations^85^ in each of the 46 tissues.

### 4.3 Identification of eVNTRs

Before the analysis of the association of VNTR genotypes and gene expression levels, we adjusted gene expression levels for each tissue in order to control for covariates of sex, population structure, and technical variations in measuring expression. For population structure, we used the top ten principal components (PCs) from a principal components analysis (PCA) on the matrix of SNP genotypes using smartpca^82^ to provide a correction for population structure. To generate the SNP genotype matrix, we used the VCF files for GTEx cohort (accession phg001219.v1) and filtered biallelic SNP sites MAF *>* 0.05 using plink^83^. To correct for non-genetic factors such as technical variations in measuring RNA expression levels (e.g batch effects, environmental variables), we applied PEER factor correction and used the top 15 factors^48^. We removed the effect of covariates by regressing them out from the RNA expression matrix of each tissue and subtracting their factor contributions and used the residuals for all eQTL association analyses.

Let *v* denote a VNTR-gene pair, *y_iv_* denote the normalized expression value of gene in *v* for individual *i* and *x_iv_* denote the genotype of the VNTR in *v* for individual *i*. Then,

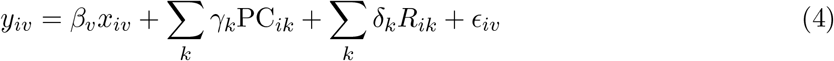

where, PC_*ik*_ denotes the strength of the *k*-th principal component, and *R_ik_* the value of the *k*-th PEER factor. We performed the association test for each VNTR-gene pair separately for each tissue type using Python statsmodels linear regression, Ordinary Least Squares (OLS)^84^, and computed a nominal p-value of the strength of association for each VNTR-gene pair.

#### Multiple Testing Correction

We used permutation tests and the BenjaminiHochberg procedure to estimate a 5% False Discovery Rate (FDR) significance cut-off for each tissue. The significance thresholds for each of the 46 tissues ranged from 10^−3^ to 3.8 × 10^−5^ (Fig. S6). Overall, 759 significant tests were observed from total of 73,609 tests in all tissues and 163 unique VNTRs passed the significance test in at least one tissue.

#### Fine-mapping of Causal Variants

To compare the strength of the VNTR association relative to proximal SNPs, we extracted all SNPs from 50kb 5’ to the transcription start, from the gene body, and up to 50kb 3’ to the end of the transcript using the GTEx variant calls. To perform a fair comparison, we used the same test and covariates for VNTRs and repeated it for each SNP by replacing the genotype to obtain the strength of association for each SNP. Then, we ranked all variants based on their association P value.

We further used a fine-mapping method, CAVIAR, as an orthogonal method to identify the causal variant for the change in gene expression level. CAVIAR is a statistical method that quantifies the probability that a variant is causal by combining association signals (i.e., summary level Z-scores) and linkage disequilibrium (LD) structure between every pair of variants^52^. We ran CAVIAR with parameter -c 1 to identify the most likely causal variant, along with the causality probability distribution for each variant site. We ranked variants based on their causality probability given by CAVIAR and called it the causality rank.

## Supplementary Figures

**Figure S1:**
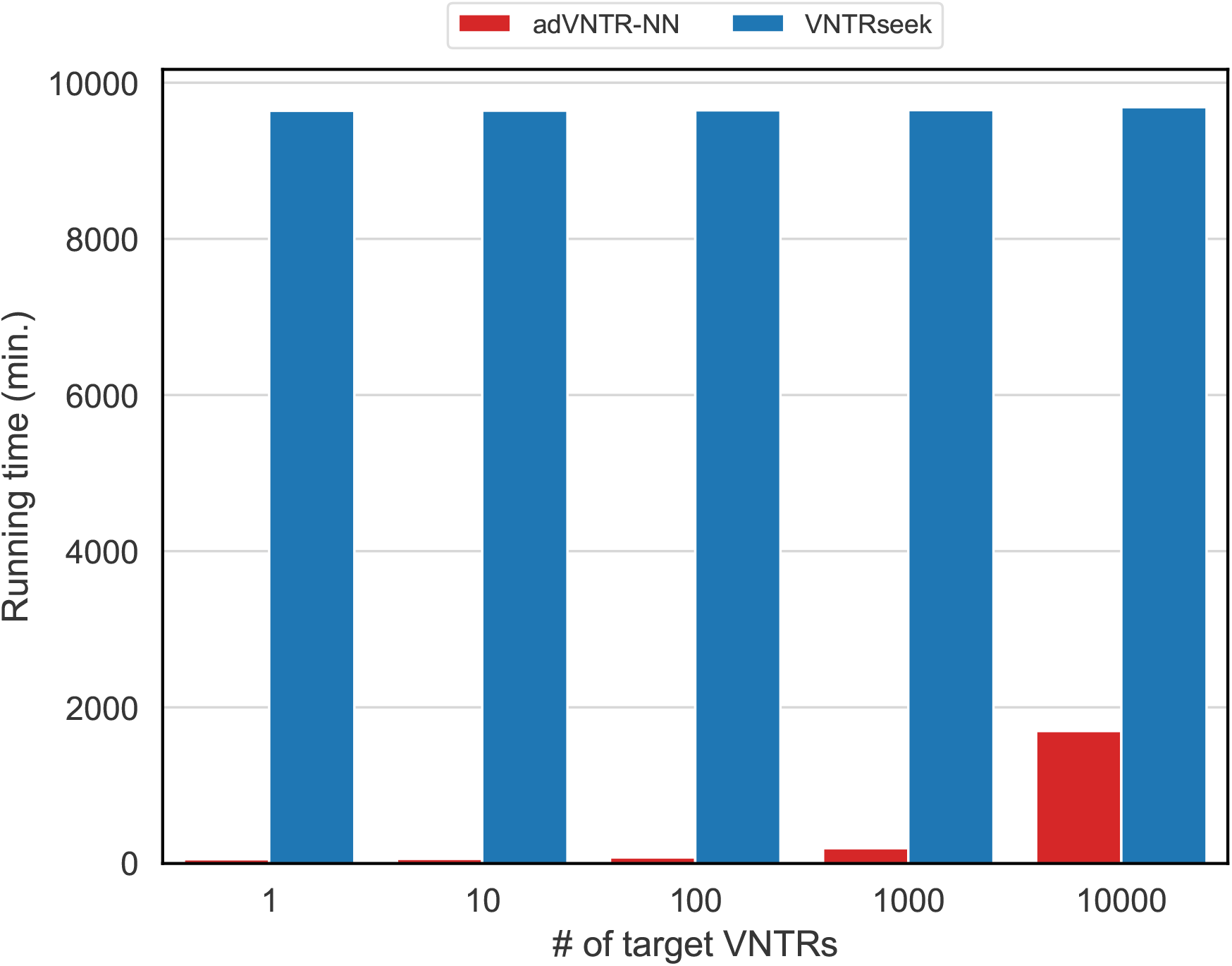
Running time comparison. Running time comparison on 1, 10, 100, 1,000, and 10,000 VNTR loci of one individual (NA24149) with 1.16 × 10^9^ reads.

**Figure S2:**
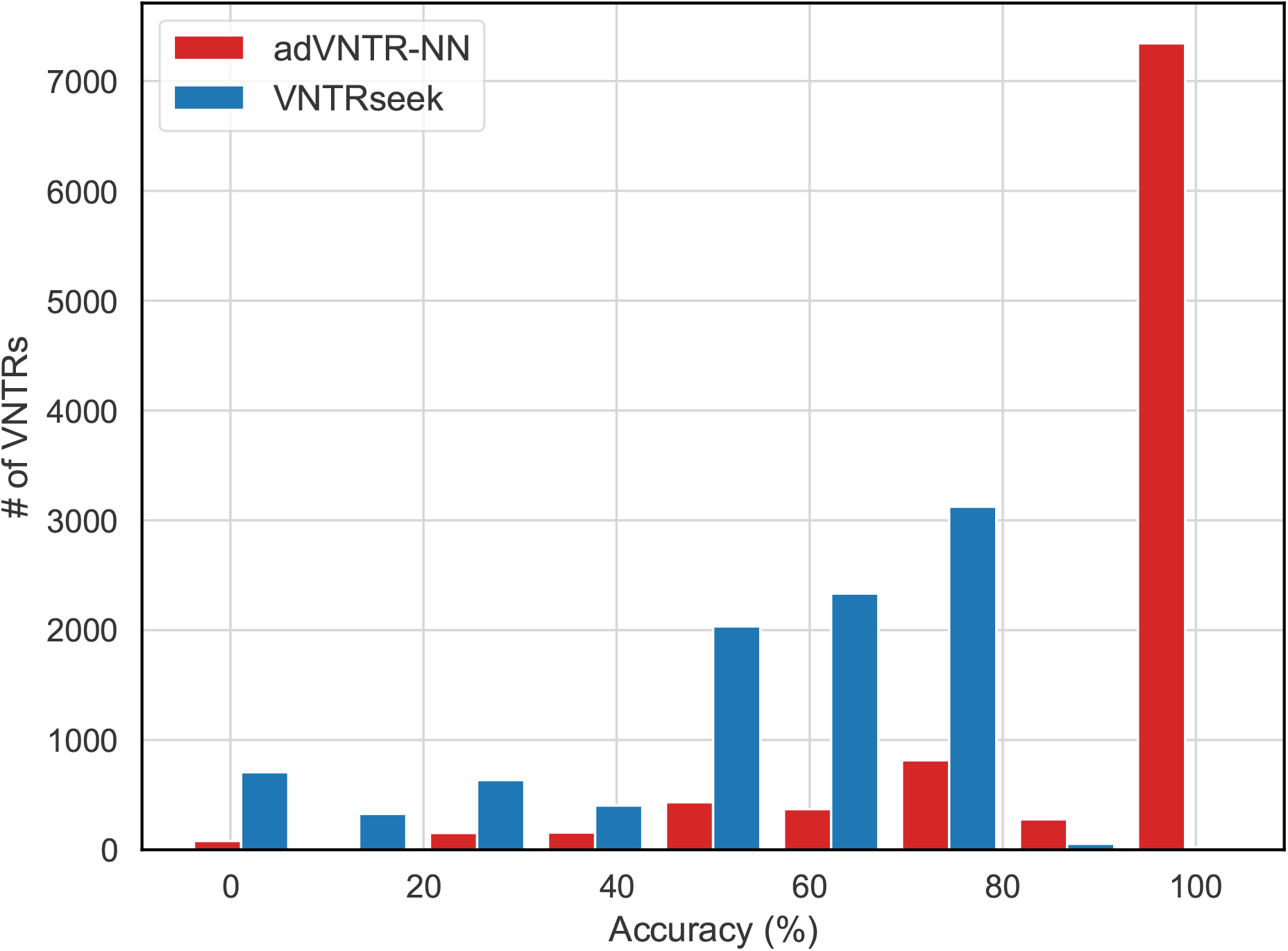
Distribution of genotyping accuracy of adVNTR and VNTRseek. The genotyping accuracy for each VNTR is defined by the # of scenarios genotyped correctly divided by # of scenarios. Six different heterozygous VNTR scenarios were tested; specifically, c/c-3, c/c-2, c/c-1, c/c+1, c/c+2, c/c+3, where c is the hg19 reference count. The number of VNTR loci modified for contraction scenarios were 9,638 (c-1), 5,078 (c-2), and 2,084 (c-3), with the reductions happening due to a requirement of at least 1 repeating copy for each VNTR allele. All expansion scenarios had 9, 638 VNTRs. adVNTR-NN had 100% accuracy in 7343 (76%) of 9638 VNTRs.

**Figure S3:**
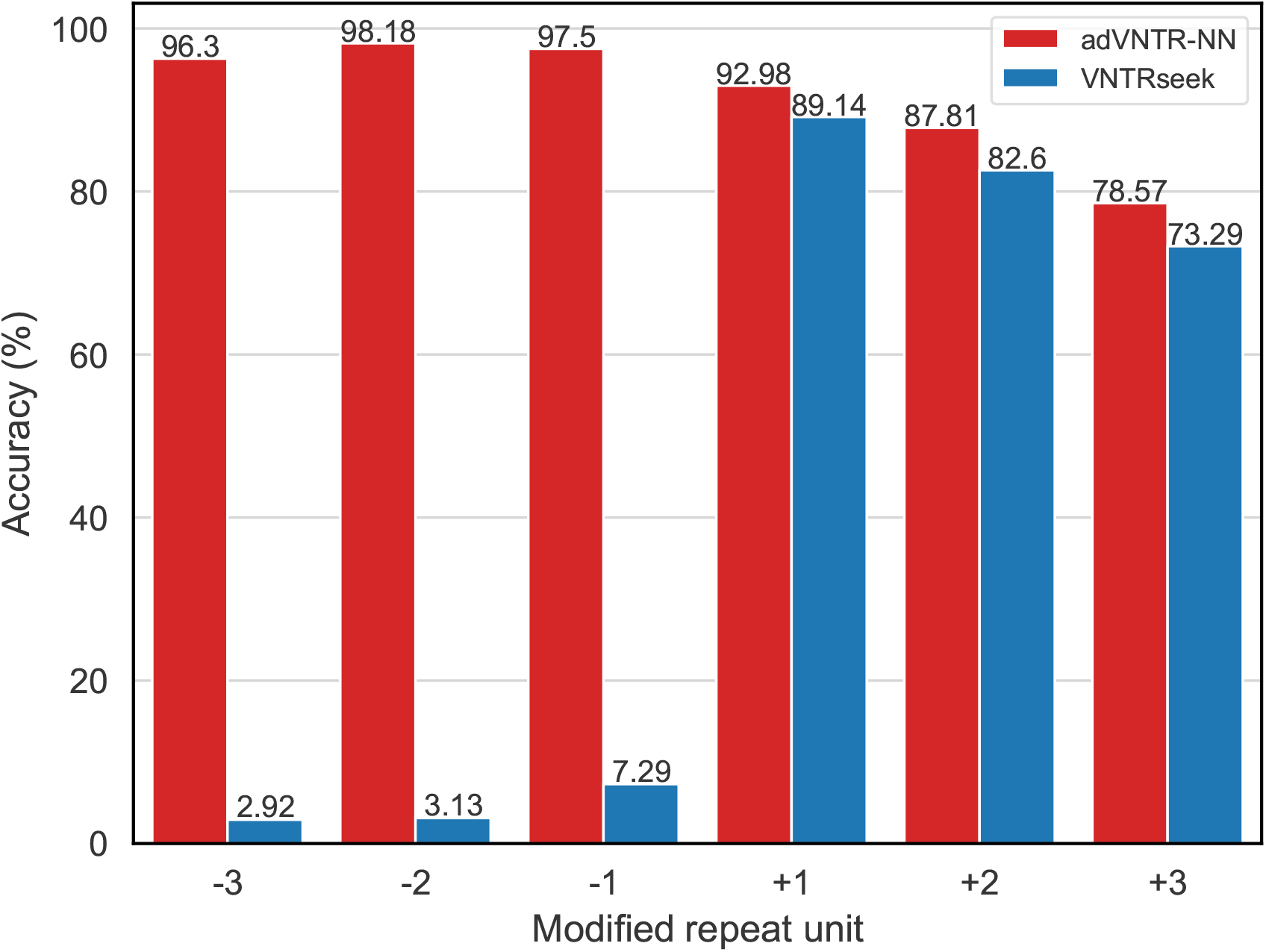
adVNTR-NN and VNTRseek accuracy for each test scenario. The genotyping accuracy for each VNTR is defined by the # of scenarios genotyped correctly divided by # of scenarios. Six different heterozygous VNTR scenarios were tested; specifically, c/c-3, c/c-2, c/c-1, c/c+1, c/c+2, c/c+3, where c is the hg19 reference count. The number of VNTR loci modified for contraction scenarios were 9,638 (c-1), 5,078 (c-2), and 2,084 (c-3), with the reductions happening due to a requirement of at least 1 repeating copy for each VNTR allele. All expansion scenarios had 9, 638 VNTRs.

**Figure S4:**
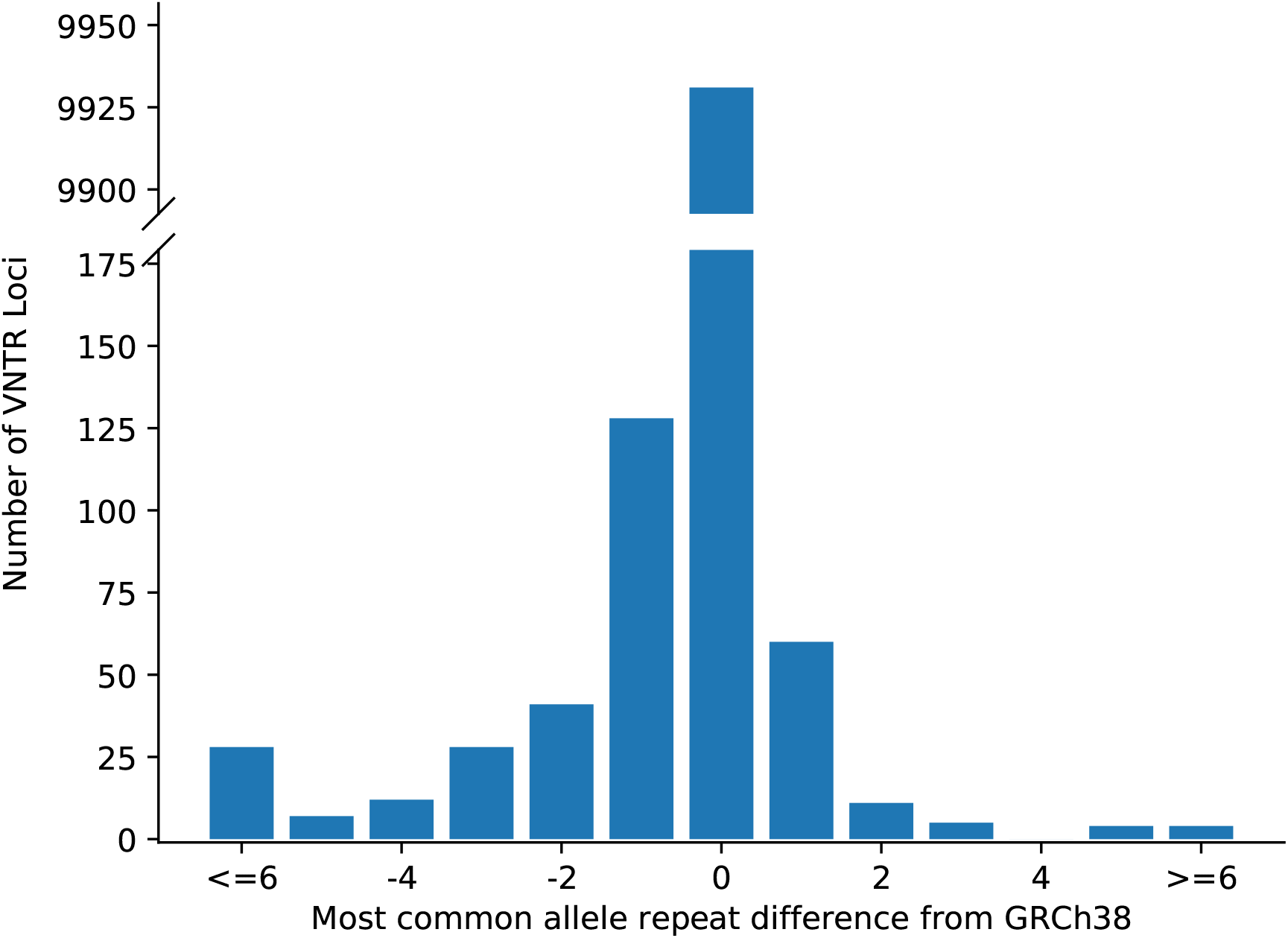
Difference in VNTR loci between donors and GRCh38. For each VNTR, the difference between the most common allele in the GTEx cohort and the GRCh38 reference repeat count was recorded. The plot shows the distribution of the differences.

**Figure S5:**
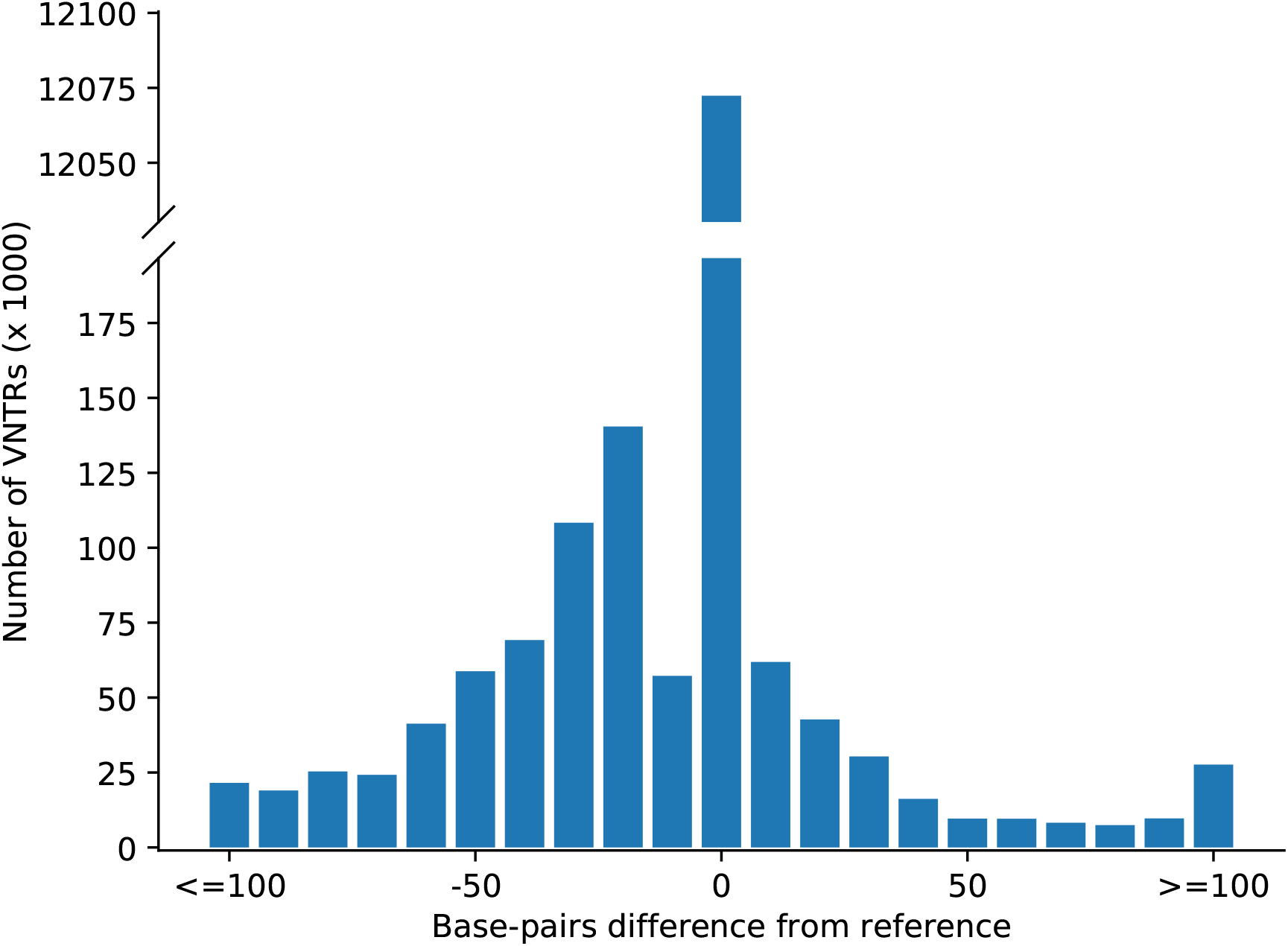
Difference in VNTR loci between donors and GRCh38. For each VNTR and each individual allele in a GTEx donor, the difference in length from the GRCh38 reference VNTR length was recorded. The plot shows a distribution of differences.

**Figure S6:**
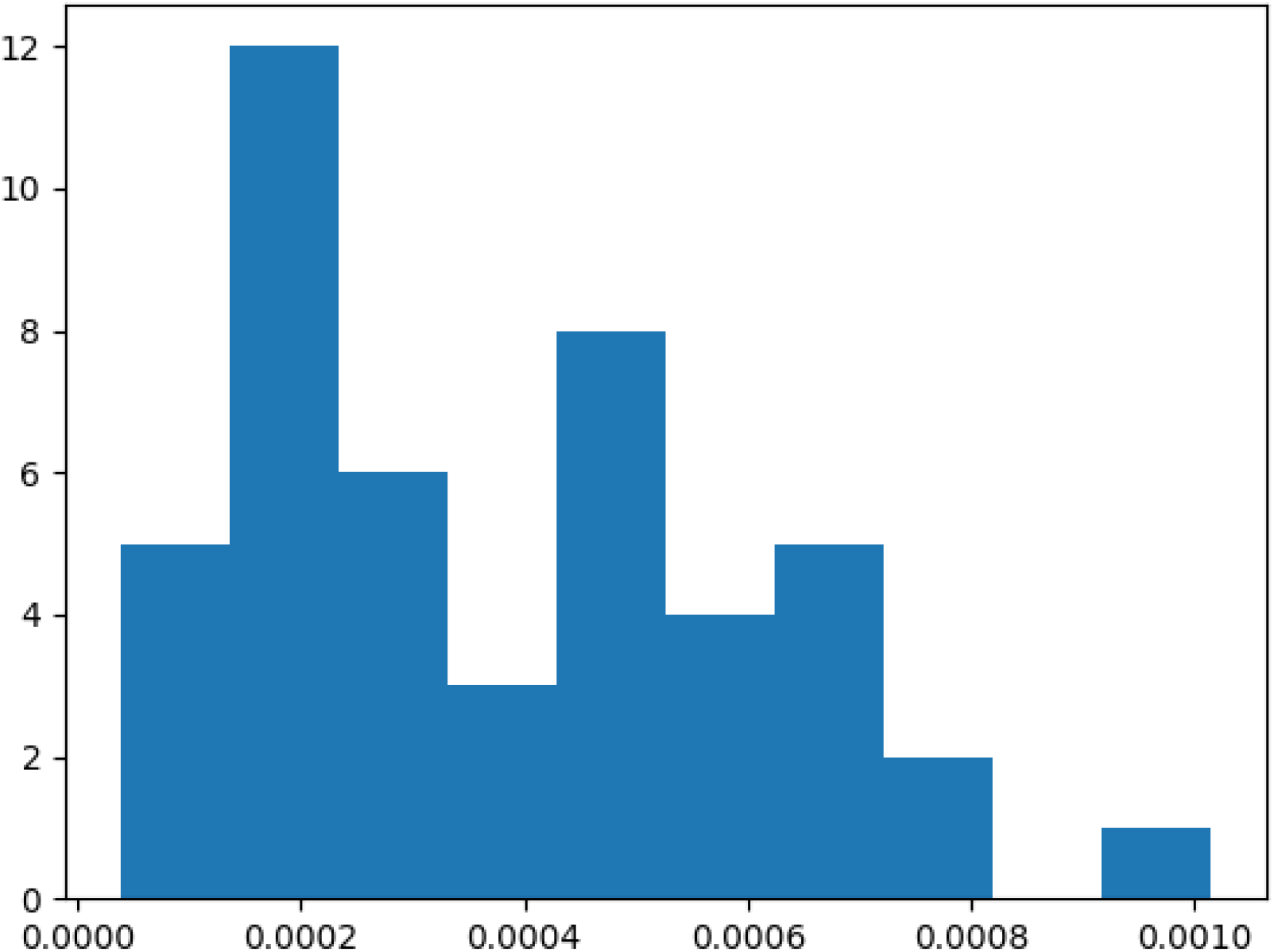
Distribution of significance thresholds for association test. Significance thresholds for each of the 46 tissues.

**Figure S7:**
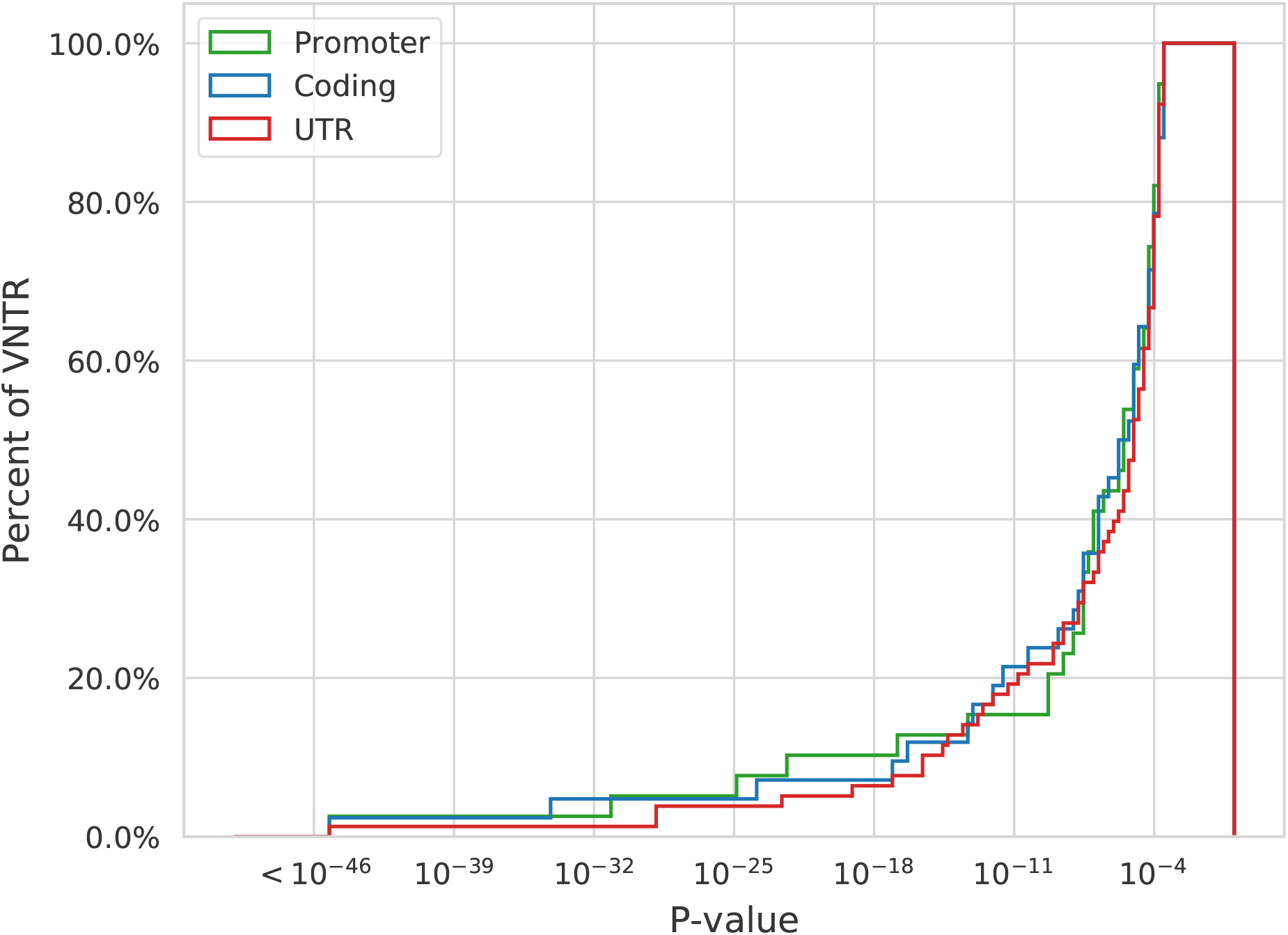
Cumulative distribution of eVNTR p-values for different classes. The plots suggest that the relative location of a genic VNTR does not significantly change the strength of association with gene expression.

**Figure S8:**
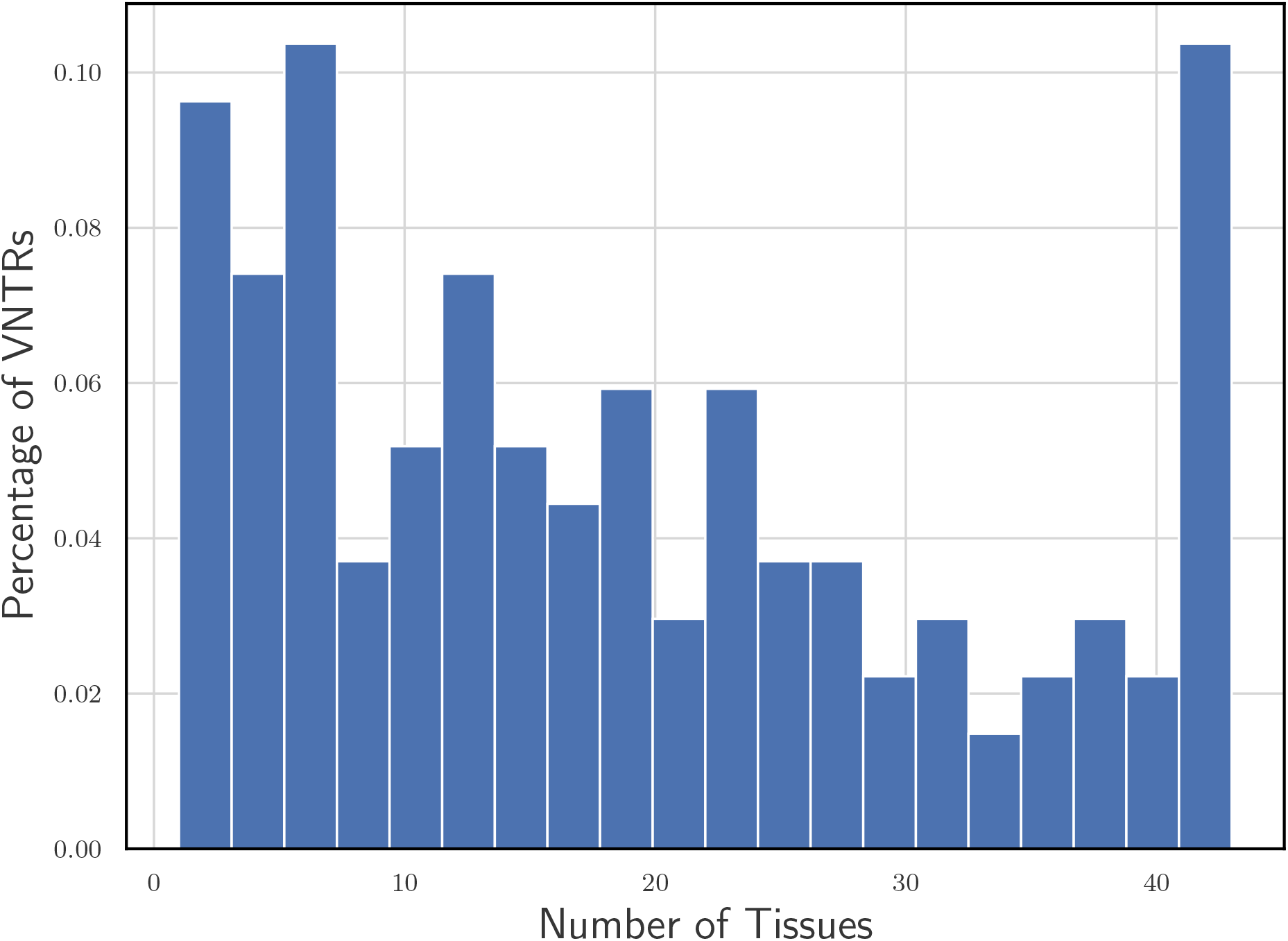
Tissue sharing of eVNTRs. The fraction of eVNTRs that are active in a specific number of tissues as determined by mash. 38% of eVNTRs were significant in at least half (23) of all tissues.

**Figure S9:**
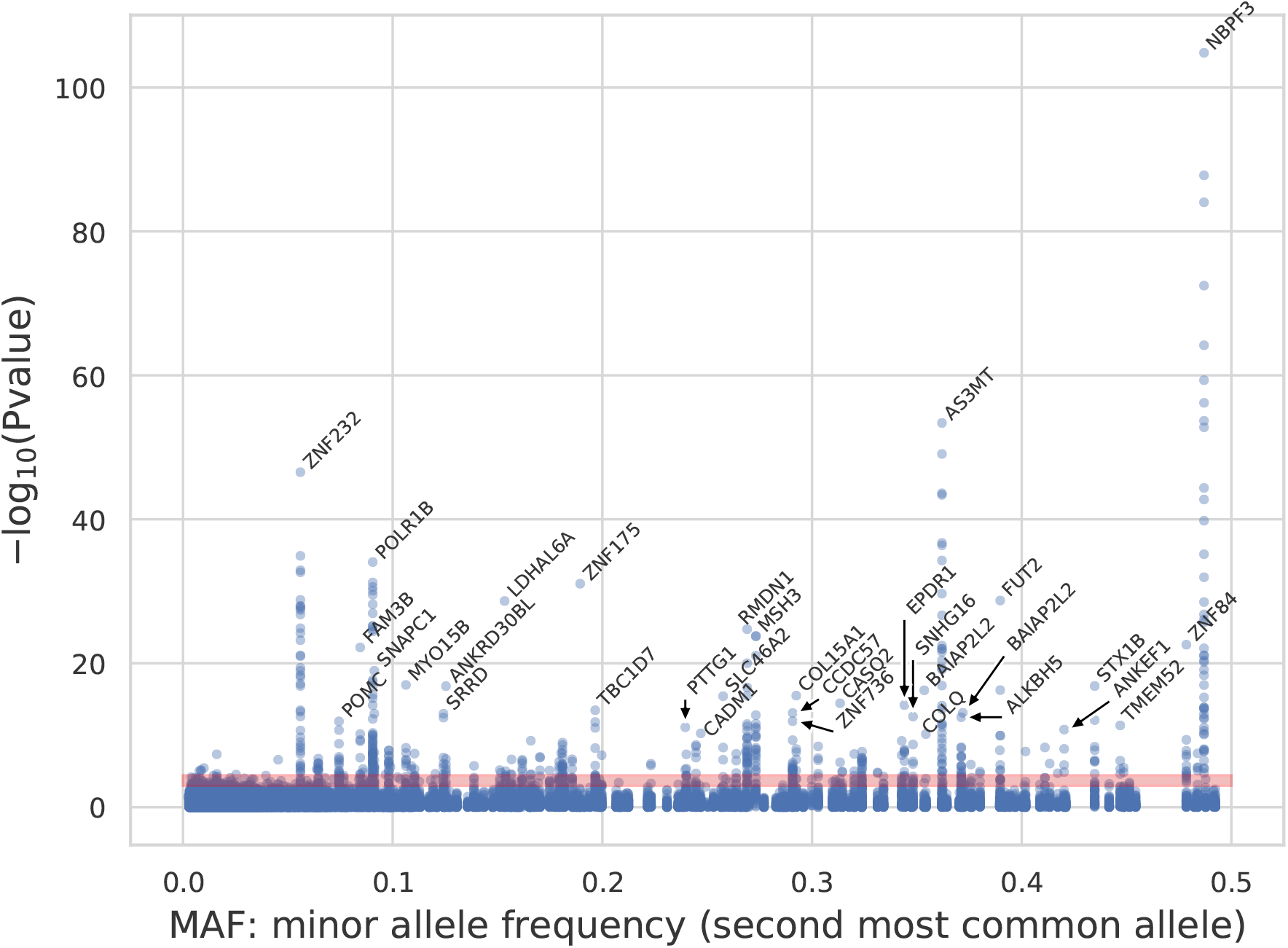
Significance of VNTR association with gene expression plotted against Minor Allele Frequency. The shaded region represents tissue specific false discovery rate cut-offs. Note that all significant tests for a single VNTR appear in a single column.

**Figure S10:**
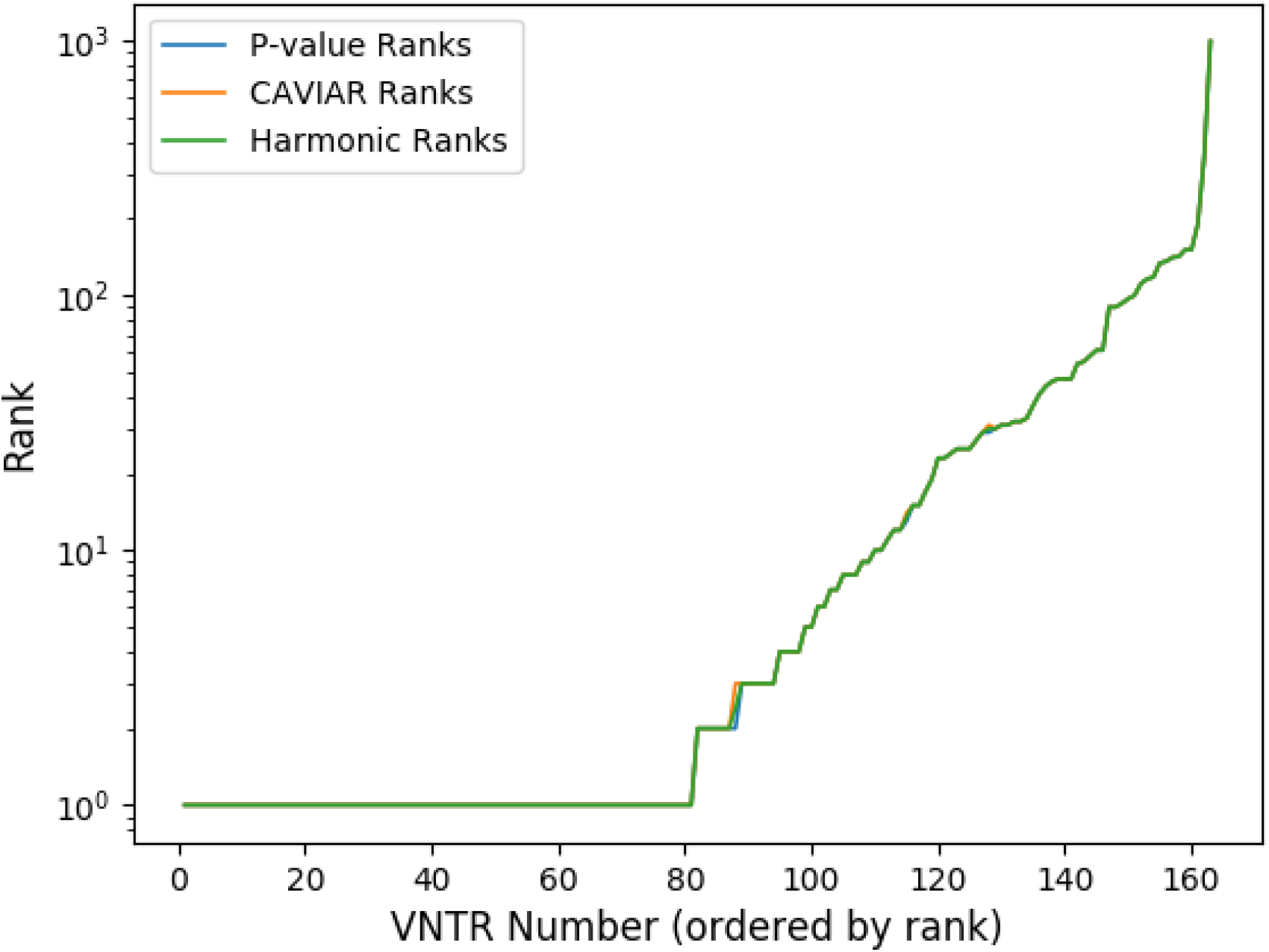
Causality rank of eVNTRs measured using strength of association (blue), CAVIAR (red), and mean harmonic rank (green). The P-value and CAVIAR based ranks coincide.

**Figure S11:**
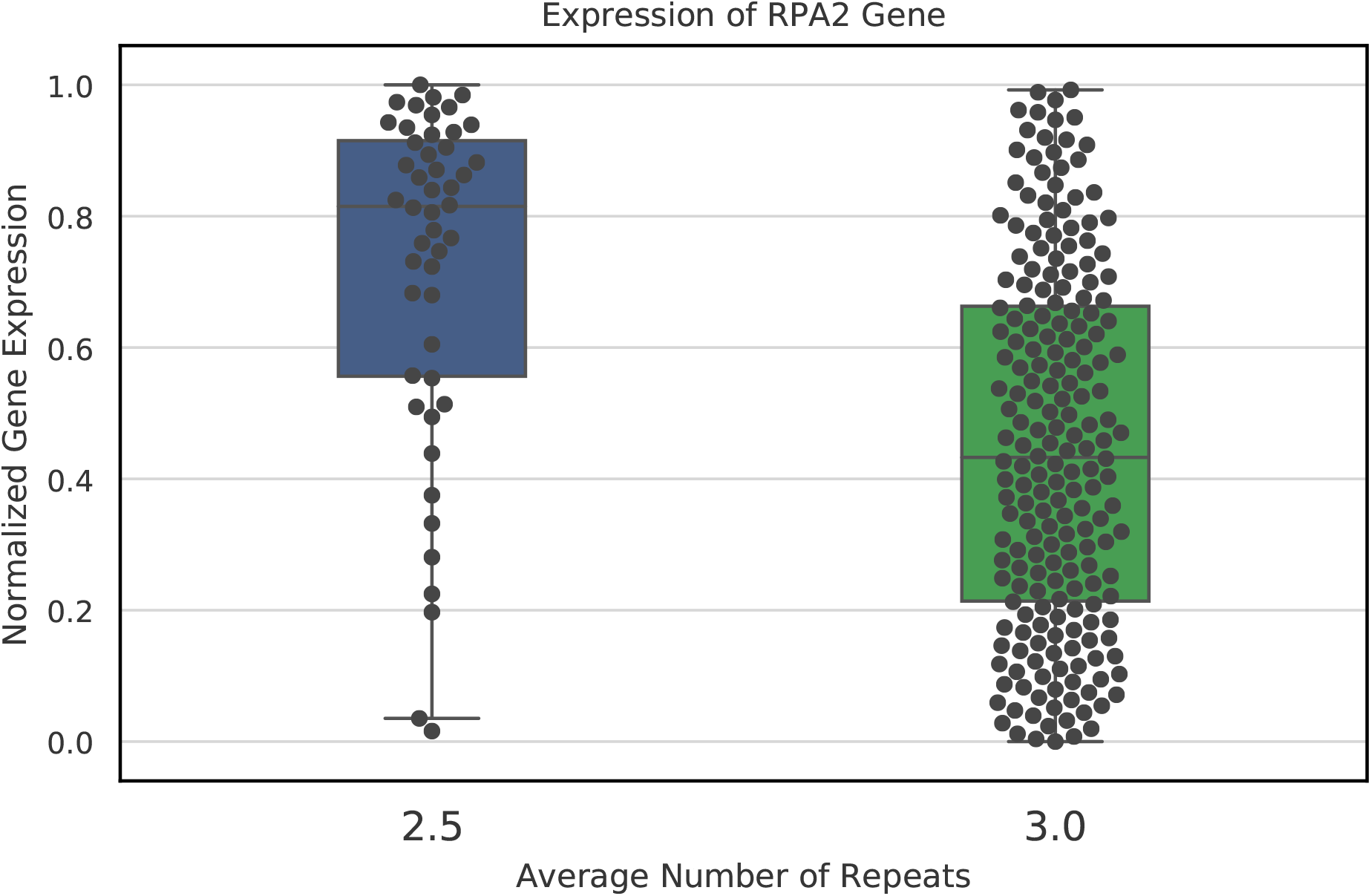
Association of RPA2 VNTR genotype with gene expression level. n=254, P-value 3.79 × 10^−25^. Increase RPA2 expression has been associated with worse survival outcomes in colon cancer^63^.

## S0.4 Neural Network Parameter Tuning

**Figure S12:**
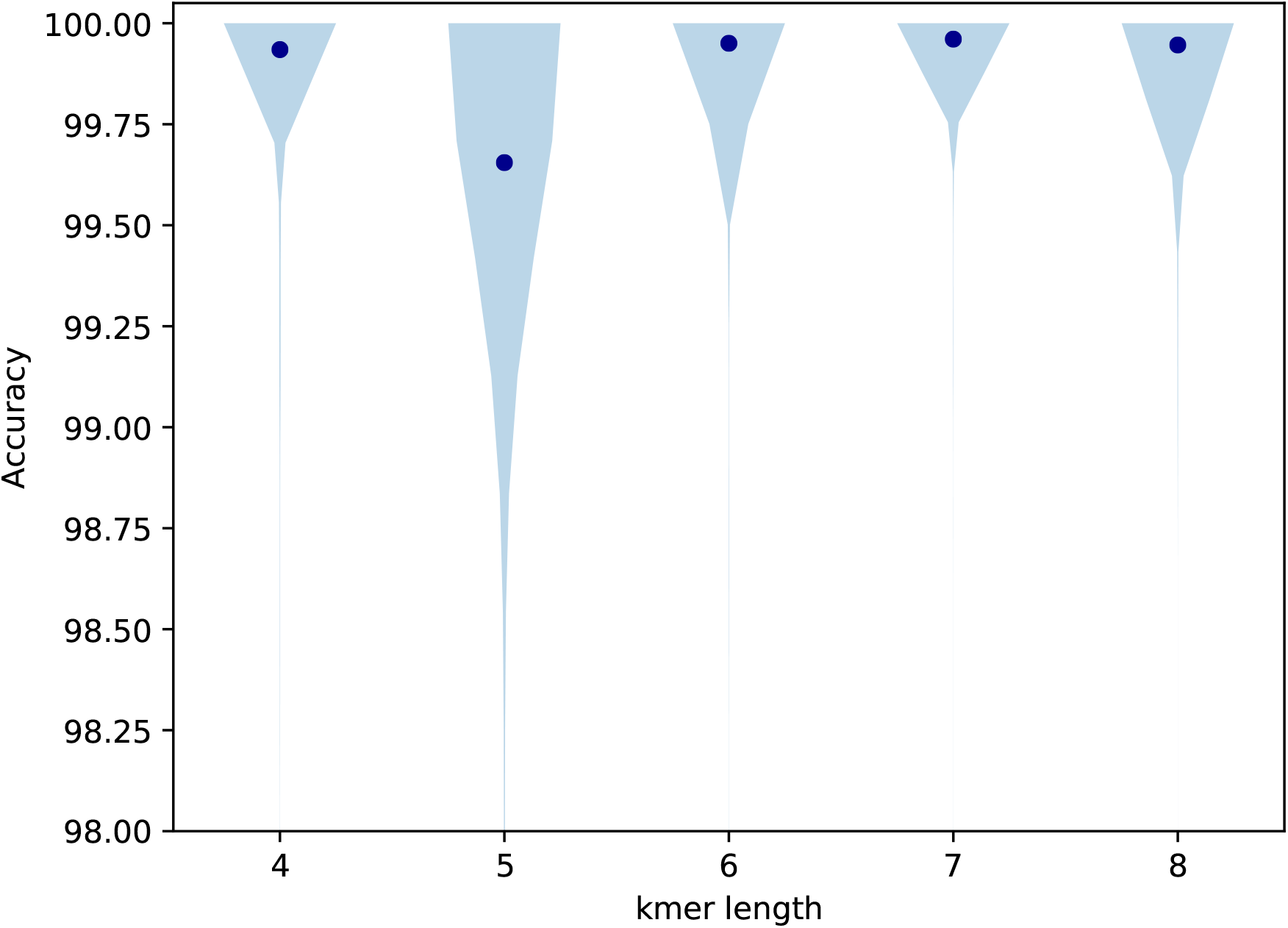
Effect of kmer length on accuracy. Performance of the neural network model on validation set for different k-mer lengths. k=6 was used for all test runs as it had the highest mean accuracy of 99.95%.

**Figure S13:**
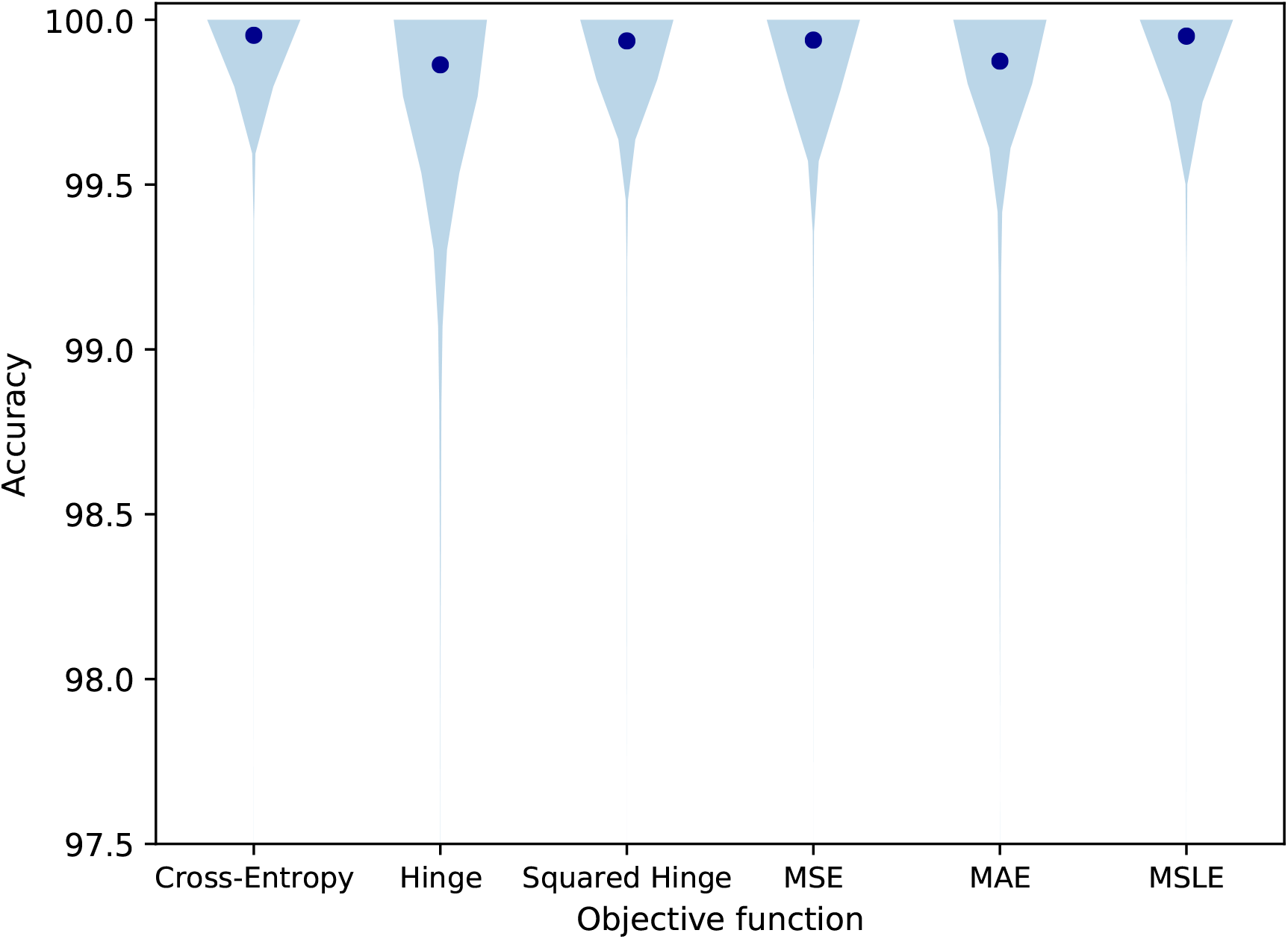
Effect of loss function on accuracy. Performance of the neural network model on validation set for different loss functions. The mean of each distribution is shown by a blue dot. Binary cross-entropy was used as the loss function for all tests.

Table S1: A list of 10,264 target VNTR loci used in this study (supplementary table s1.xlsx)

